# A Conserved Mammalian Hippocampal Navigation Motif Reconfigured for Primate Vision

**DOI:** 10.1101/2025.07.01.662661

**Authors:** Carlos Otero, Diego B. Piza, Ehsan Aboutorabi, Julio C. Martinez-Trujillo, Jorge Riera

## Abstract

Spatial navigation requires the brain to continuously sample the external world while evaluating internal representations of space. In rodents, this process unfolds through alternating periods of locomotion and pauses. During pauses, hippocampal sharp-wave ripples (SWRs) rates increase, likely reflecting the broadcast of spatial memories that guide navigation. Whether similar dynamics govern navigation in primates remains unknown. Here, we recorded hippocampal activity in freely moving marmosets navigating a 3D maze. Like rodents, marmosets alternated between locomotion and pauses. However, pauses were long and showed an increase in SWR rates relative to locomotion. SWRs were most prominent when animals maintained stable head orientations toward rewarded locations and were reduced during rapid exploratory head movements. SWR rates further increased when spatial memories were used to guide navigation. Our findings reveal a phylogenetically conserved motif linking behavioral states during spatial navigation to hippocampal SWR dynamics across mammals and show how primate visual specializations have adapted this motif to support vision-guided navigation.

**Significance Statement:** Our results reveal a phylogenetically conserved hippocampal navigation motif that has persisted despite major evolutionary changes in mammalian sensory ecology. Across species, navigation alternates between external exploration and internal evaluation, with SWRs marking periods of memory-guided computation. However, primate evolution reshaped the behavioral expression of this motif by coupling it to active visual sampling, gaze control, and foveal inspection of landmarks. Thus, evolution appears to have preserved a core hippocampal algorithm for navigation while adapting its sensory inputs and behavioral context to the demands of diurnal, vision-guided life.

## Introduction

Spatial navigation requires integrating incoming sensory information with stored representations of space, a process thought to rely on computations performed by hippocampal circuits **^[1–3]^**. Studies in nocturnal rodents such as mice and rats show that navigation does not proceed continuously but unfolds through alternation between locomotion toward goals and brief pauses **^[4–7]^**. During locomotion, sensory sampling is driven by rhythmic scanning via olfaction and whisking, as well as by vestibular and proprioceptive inputs that aid path integration. During pauses, rodents explore the environment via less rhythmic behaviors, such as rearing, that allow memory-guided evaluation and updating of spatial position **^[8–9]^**. Thus, rodent navigation follows a structured motif in which locomotion-pause alternations support multimodal sensory sampling and interactions with stored cognitive maps of space, thereby contributing to route updating **^[10]^**. Whether this motif is conserved in diurnal primates, which evolved high-acuity color vision and foveal specialization, and whose active sensing during navigation relies predominantly on visual gaze exploration **^[11]^**remains unknown.

In the rodent hippocampus (HPC), this behavioral alternation is mirrored by transitions between theta rhythms, which dominate during locomotion, and sharp-wave ripples (SWRs), which dominate during pauses **^[4–8]^**. Theta rhythm has been linked to online sensory sampling and navigation, while SWRs are associated with the internally initiated broadcast of hippocampal representations to distributed brain networks **^[12–14]^**. However, in other mammals such as primates, theta rhythms are rare, and SWR rates have not been explored during navigation **^[11]^**. Here, we will focus on SWRs, synchronous neurophysiological events occurring in the HPC characterized by the co-occurrence of low-frequency sharp waves (SPW) and high-frequency Ripples, which are widely implicated in memory formation, consolidation, and recall **^[5,15–17]^**. SWRs have been observed across species, including rodents **^[18–19]^**, bats **^[20]^**, reptiles **^[21]^**, and birds **^[22]^**. In freely navigating rodents, SWRs increase in frequency during pauses in locomotion and with memory demands **^[5]^**. It is thought that during pauses, externally driven sensory input is reduced, while internally driven cognitive processing increases. SWRs emerge at behaviorally relevant moments, such as at reward sites or decision points, and are associated with replay or preplay of place-cell sequences and the evaluation of alternative paths. Disrupting SWRs during these periods impairs learning and decision-making, underscoring their role in internal computation rather than behavioral disengagement **^[5,23–26]^**.

In primates, the relationship between SWRs and spatial navigation remains largely unexplored **^[27–28]^**. However, over the last decades, advances in intracranial recording techniques and neurosurgical approaches have opened new avenues for studying hippocampal dynamics in primates **^[11,14,29–36]^**. Studies in macaques and human patients have revealed a relationship between SWR occurrence and visual exploration, particularly during saccadic eye movements **^[33–35,37–39]^**. However, because these experiments were performed under head-restrained conditions, they preclude direct comparison with navigation studies in freely moving rodents. The common marmoset (Callithrix jacchus), a small New World primate, provides a unique opportunity to fill this gap in the study of hippocampal function during spatial navigation **^[40]^**.

A recent study reported that freely moving marmosets also alternate locomotion with pauses during navigation **^[11]^**, suggesting that the behavioral motif described in rodents may extend to primates. The same study has shown that during pauses, marmosets make rapid exploratory head movements that orient gaze and the fovea toward visual landmarks. These gaze-orienting head movements are absent in rats and may play a fundamental role in shaping view-cell selectivity in the hippocampus **^[11,31,41–42]^**. One may argue that marmoset visual exploratory pauses resemble rearing behavior in rodents, since in both scenarios animals engage in active sensory sampling of the environment **^[10–11]^**. We hypothesize that, in freely moving marmosets, hippocampal SWRs that broadcast stored spatial representations preferentially occur during pauses, and increases in SWR rates mark a transition from externally driven sensory sampling during locomotion to internally driven evaluation. We further hypothesize that SWR rates further increase when spatial memories must be used to guide navigation.

We tested these hypotheses by wirelessly recording hippocampal local field potentials (LFPs) from freely moving marmosets performing spatial navigation tasks. We implemented and validated a SWR detection framework that combines dual-band filtering, coincident sharp wave (SPW) and Ripple criteria, and observer-verified inspection, enabling the extraction of detailed spectral and temporal features **^[43–44]^**. Recordings were obtained from animals performing both simple foraging and memory-guided alternation tasks in a 3D maze equipped with synchronized motion tracking. Each animal was implanted with a chronic microelectrode array targeting hippocampal regions CA1 and CA3, allowing us to relate neural dynamics to precise measurements of head and body movements. We found that SWR rates were higher during pauses than during locomotion. During pauses, SWR rates were highest when animals oriented their heads (gaze) toward behaviorally relevant, rewarded locations and decreased when their heads were moving. Importantly, SWR rates increased further when animals used spatial memory to guide navigation.

## Results

We recorded neural activity in the hippocampus of three freely moving marmosets implanted with microelectrode arrays (brush MEA, Microprobe Inc., MD) while tracking their position using a motion-capture system with reflective markers (Optitrack, Natural Point Inc., OR). We recorded LFPs from subfields CA1 and CA3, and applied a SWR detection algorithm (see Methods). We detected a total of 19,481 SWRs recorded across 49 sessions in CA1 (Subject C, left HPC, 15 sessions, n=2960 events) and CA3 (Subject PB, right HPC, 9 sessions, n=3812 events) (Subject L, right HPC, 25 sessions, n=12,709 events) (see **Supplementary Table 2 & Table 3** in Methods).

During the sessions, animals navigated the 3D maze to obtain rewards at ports located on the maze walls (**Fig. 1A**). We tested the animals during two different tasks: 1) a foraging task, in which a single reward site was cued with a flashing LED in pseudo-random order, and the animal received a liquid reward after entering the active reward zone (29 sessions); and 2) a memory alternation task, in which rewards alternated between selected sites, requiring the animal to learn and recall the sites to retrieve reward correctly (30 sessions) (**Fig. 1B**).

**Figure 1.**
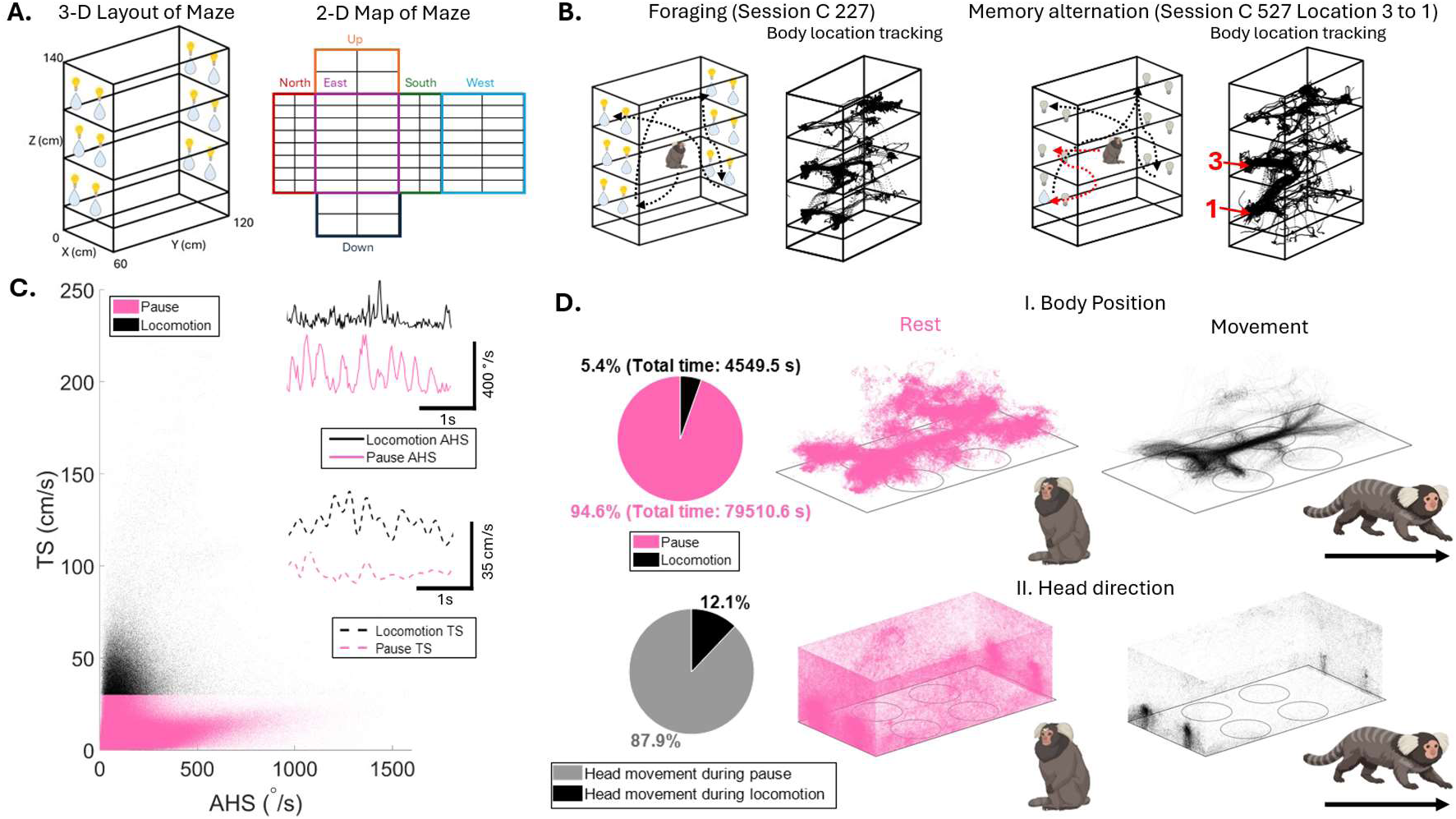
Experimental set-up and marmoset movement tracking **A.** Layout of the 3D maze is shown. To the right, the maze walls, including the ceiling and bottom-most floor, are unfolded to project the subregions of the maze. **B.** Example body position tracking during a foraging (C 227, left) and memory alternation session (C 527, right). Each contains a cartoon showing the typical movement pattern for each case, illustrating the randomness of foraging compared to the sequence alternation in the memory task (Dashed red trajectories represent rewarded sequence in the memory alternation task).**C.** Scatter plot of the TS (cm/s) versus AHS (°/s). Body pauses data shown in pink and locomotion in black. Example traces for each epoch are also shown; solid lines represent the AHS and dashed lines the TS, with scale bars on the bottom right. The speed threshold was set to 30 cm/s (See Methods). **D.** Pie charts show the proportion of time subjects spent in pauses and locomotion during the recordings (along with the total duration(s)). The right panels show two illustrations of the second floor of the maze with: **I)** traces of body position during pauses (top) and head direction during pauses (bottom) (49 sessions), and **II)** traces of body position during locomotion (top) and head direction (bottom) (49 sessions).

We computed translational speed (TS) and head angular speed (AHS) from position signals across all sessions (**Fig. 1C**). We identified two states, corresponding to pauses (TS < 30 cm/s) and locomotion (TS > 30 cm/s, locomotion) (pink area in **Fig. 1C**). The speed threshold (30cm/s) was based on a travel distance of approximately one marmoset body length (∼25-30cm) in a second **^[45]^**. Interestingly, animals spent 94.6% of the time (79,510.6 s) in the pause state, and 5.4% of the time locomoting through the maze (4,549.5 s). Moreover, 87.9% of head movements occurred during pauses (**Fig. 1D**, see Methods for thresholding criteria). These dynamics contrast with those in rodents, which display larger head movements during full-body turns and smaller, slower head movements during body pauses **^[11]^**. Marmosets exhibit rapid head movements primarily during pauses in body movement (see **Fig. 1D** pie charts and **Supplementary Fig. 1**). We attribute this behavior to their ability to detect and locate reward sites at a distance using high-resolution color-stereo-vision (exploration without visitation), thereby avoiding the energy expenditure of continuously exploring the environment during locomotion via whisker-based somatosensing and olfaction **^[11]^**.

We further isolated the body position in the 3D maze coordinates (see one floor example in **Fig. 1D I**) and head direction by ‘plotting’ the line of view on the maze walls surface (**Fig. 1D II**) during pauses (pink and left columns), and during locomotion (black dots right column). In both cases, the position tends to align with the areas of the maze where the animals navigated (trajectories), suggesting that the animals paused along the same navigation paths that follow the maze geometry (i.e., they avoided the apertures in the middle when traveling along a floor path). The head pointing direction, which we consider an estimate of gaze direction, covered the entire maze walls; however, it concentrated on the reward locations, indicating that animals were actively sampling those locations, likely to reorient themselves, during both pauses and locomotion.

### SWR detection and spatiotemporal features

To explore hippocampal SWR rates across tasks and during locomotion and pauses, we developed a detection method that utilized spectral power thresholding in the Ripple and SPW bands to find periods of increased power in both these frequency ranges **^[43]^** (**Fig. 2A**). We also used other criteria such as minimum Ripple duration, transient artifact rejection, and power-line noise removal (See Methods for details). To validate our detection, we evaluated it on experimentally recorded LFP segments in which three expert annotators independently labeled SWRs. Only marked intervals that were overlapped by all three experts were considered ground-truth events. A detected event was considered a true positive if its detected onset and offset each fell within ±25 ms of the corresponding ground-truth onset and offset (**Fig. 2B**), while time samples of no SWR activity were marked as true negatives. All detectors included an adjustable parameter for the Ripple power z-score threshold, so that parameter was changed across the detection sweeps. For this study, a setting of 3 standard deviations from the mean of the z-score for Ripple power was used (See methods).

**Figure 2.**
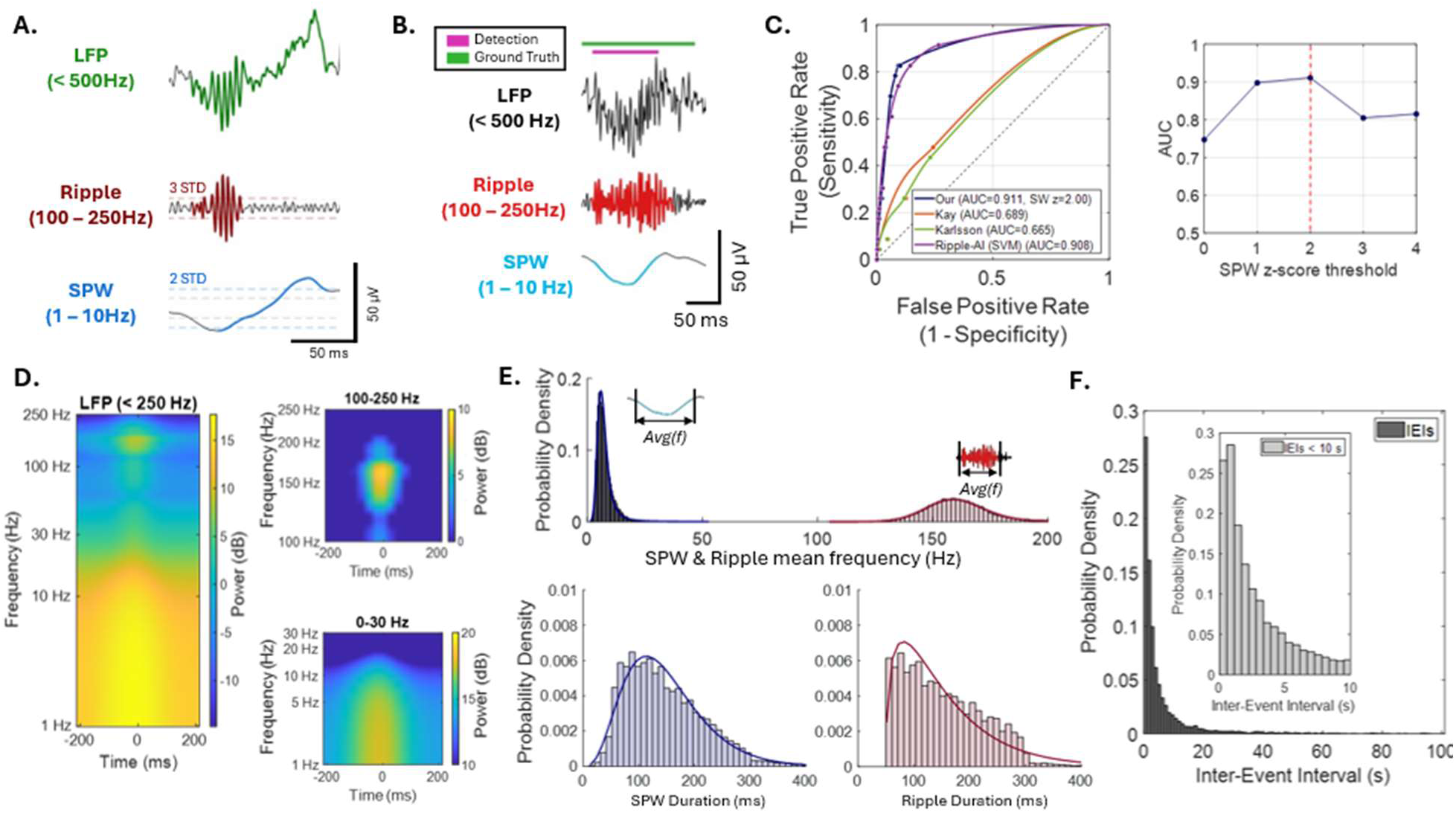
Validation of detection and extraction of SWR features **A.** Classical SWR example with the LFP signal (green) on the top, Ripple (red, 100-250 Hz) in the middle, and SPW (blue, 1-10 Hz) on the bottom row. The Ripple and SPW components are shown with colored dashed lines to represent the threshold of event detection (3 STD for Ripple, 2 STD for SPW) for the start of the component and grey dashed lines (1 STD for both components) to show the lower threshold for the end of the component. Scale bars on the bottom right show voltage versus time (See Methods). **B.** An example of a ground truth SWR identified by the three experts. The top row is the LFP, middle is the Ripple (red), and bottom is the SPW (blue) (same frequency bands as Fig. 2A). The solid bars on the top represent the interval that was classified as an event, green for the ground truth overlap between the experts, purple for the detection method applied. Scale bars on the bottom right indicate voltage vs time units. **C.** Validation of our SWR detection method compared with three established Ripple detectors **^[13,44,46]^**. Left: ROC curves showing our method (blue) versus Kay (orange), Karlsson (green), and Ripple-AI SVM (purple), computed by varying the Ripple threshold in each detector. ROC area shown in legend. Right: AUC for our method across a range of SPW Z-score thresholds. The vertical red dashed line represents the upper threshold for the SPW detection used in our study. **D.** Grand-average LFP spectrogram across SWRs (n = 19,481 events), using a 400-ms window centered on the Ripple-peak. The right panels show zoomed-in views of the 100–250 Hz (top) and 1–30 Hz (bottom) ranges. Color-bar ranges differ between panels. **E.** Probability density functions of SWR features (n = 19,481 events). First row: shows the mean frequency (Hz) of the SPW (blue) and Ripple (red) along with their stable fits; Second row: the duration (ms) of the SPW (blue) on the left and Ripple (red) on the right alongside the gamma fits of each. (See Methods for equations of distribution fittings). Illustrations in the mean-frequency panels indicate that this measure reflects the average instantaneous frequency of each component. **F.** Inter-event Intervals (seconds) of the SWRs (n = 13,579 IEIs) across 49 sessions. The histograms show the IEI distributions from 0 – 100 s (black, bins 1 s wide) with an inset of the IEIs between 1 – 10 s (gray, bins 560 ms wide).

Using this observer-validated dataset, our method outperformed three other previously used approaches (AUC (area under curve) = 0.911 for our method vs. 0.689 for Kay **^[46]^**, 0.665 for Karlsson **^[13]^**, and 0.908 for Ripple-AI **^[44]^**), further confirming its reliability for marmoset SWR detection in our recordings (**Fig. 2C**, left). The incorporation of the SPW, as well as differences in the recording electrodes used across studies, may help explain why our method and Ripple-AI may perform better than the other two methods (See ref. Liu et al. **^[43]^** for a review). As few studies have reported using the SPW component for SWR detection, we used this AUC to validate our settings: a z-score threshold of 2 for SPW-band power (**Fig. 2C**, right). Overall, we used 3 STD of the ripple power z-score and 2 STD of the SPW power z-score for SWR detection.

The grand-average spectrogram of the 19,481 detected SWRs, centered on the peak amplitude of the Ripple component, shows a concentration of spectral power at the Ripple and SPW frequencies (**Fig. 2D**). The Ripple frequency band shows a distinct maximum centered on the peak Ripple amplitude, spanning approximately 50 ms. We also observe a 25% increase in power in the SPW frequency band over approximately 200 ms. The average power and standard deviation across different frequency ranges and over the entire 400 ms time window were as follows: Full Frequency Range (1 – 250 Hz): −1.64 dB (STD: 6.41 dB), High Frequency Range (100 – 250 Hz): −4.07 dB (STD: 5.33 dB), and Low Frequency Range (1 – 30 Hz): 9.90 dB (STD: 4.20 dB).

We decomposed each SWR into SPW and Ripple components and independently quantified the duration and mean frequency of each. We also performed distribution-fitting to estimate parameters for both the stable and gamma probability density functions (PDFs). The SPW duration PDF, fitted using a gamma distribution, had parameters a=4.31 and b=34.22, with a mean duration of 147.50 ms, a median of 134.00 ms, and a standard deviation of 72.87 ms. The Ripple component, on the other hand, displayed unique temporal characteristics modeled using a gamma distribution with parameters a=1.91 and b=53.64. The Ripple had a mean duration of 152.55 ms, a median of 140.00 ms, and a standard deviation of 74.16 ms. These results demonstrate variability in the duration of the SPW and Ripple components within the explored frequency ranges (**Fig. 2E**). The gamma distribution’s parameter ‘a’ controls skewness, with lower values indicating greater skewness, while the scale parameter ‘b’ determines the spread, with larger values reflecting a wider distribution. Compared to SPWs, the Ripple components exhibited greater skewness (lower a) and a broader spread (higher b), indicating more variable and asymmetric duration distributions.

The SPW and Ripple components also exhibited distinct frequency characteristics. The SPW mean-frequency probability distribution function (PDF), fitted with a stable distribution, was characterized by parameters α=1.41, β=1.00, γ=1.53, and δ=6.21, with a mean frequency of 7.47 Hz, a median of 6.68 Hz (theta band), and a STD of 6.21 Hz. In contrast, the Ripple component exhibited higher frequencies, with its mean frequency also well fit by a stable distribution with parameters α=1.83, β=0.84, γ=9.03, and δ=159.22. The Ripple mean frequency was 160.87 Hz, with a median of 140 Hz (above the high-gamma band) and a STD of 14.09 Hz (**Fig. 2E**). The parameters capture different aspects of the stable distribution: the stability parameter (α) controls the tail thickness, with higher values indicating narrower tails; the skewness parameter (β) governs asymmetry, where values closer to zero reflect reduced skewness; the scale parameter (γ) determines the spread of the distribution; and the location parameter (δ) sets the central tendency. Compared to SWs, the Ripple components exhibited reduced skewness (β) and a greater scale (γ), consistent with a less skewed but more broadly distributed mean frequency profile.

We further explored inter-event intervals (IEIs), defined as the time between consecutive unique SWR events. Therefore, unique events were defined by collapsing overlapping SWR detections across channels, retaining only one event per overlapping group. This same unique-event definition was used for both SWR rate and IEI calculations, such that SWR rates were computed from the number of unique SWR events per recording duration. Across 49 sessions, 19,481 channel-level SWR detections were reduced to 13,628 unique SWR events, indicating that 5,853 detections (30.04%) reflected overlapping detections across channels rather than single-channel SWRs. Because the first unique event in each session lacked a preceding event to calculate an interval, these unique events yielded 13,579 IEIs across the three subjects. The mean IEI was 10.11 s, with a median of 2.34 s and a standard deviation of 39.82 s. Notably, 11,235 IEIs (82.74%) were under 10 s. These findings indicate that SWRs occurred at irregular intervals, with a bias toward shorter intervals, consistent with burst-like SWRs.

### Differences in SWR rates between foraging and memory alternation

To determine whether memory demands influence SWR dynamics during navigation, we compared SWR rates during foraging and memory-guided alternation (**Fig. 3A**). Two of our subjects (C and L) were tested in both tasks, whereas Subject PB was tested only in the foraging task. We pooled data from the 3 subjects (C, L, and PB) during foraging and from two subjects (C and L) during memory alternation. SWR rates were significantly higher during memory alternation than during foraging (one-tailed Wilcoxon rank-sum test on SWR rates, p = 7.63e-3; **Fig. 3B**, **Supplementary Table 1**). Consistent with this result, inter-event intervals (IEIs) were significantly shorter during memory alternation than during foraging (one-tailed Wilcoxon rank-sum test, p = 2.35e-32; **Fig. 3C**, **Supplementary Table 1**). We next separated each session into locomotion and pause periods (See Methods) and computed corresponding SWR rates. The results were the same when excluding subject PB (**Supplementary Fig. 3)**.

**Figure 3.**
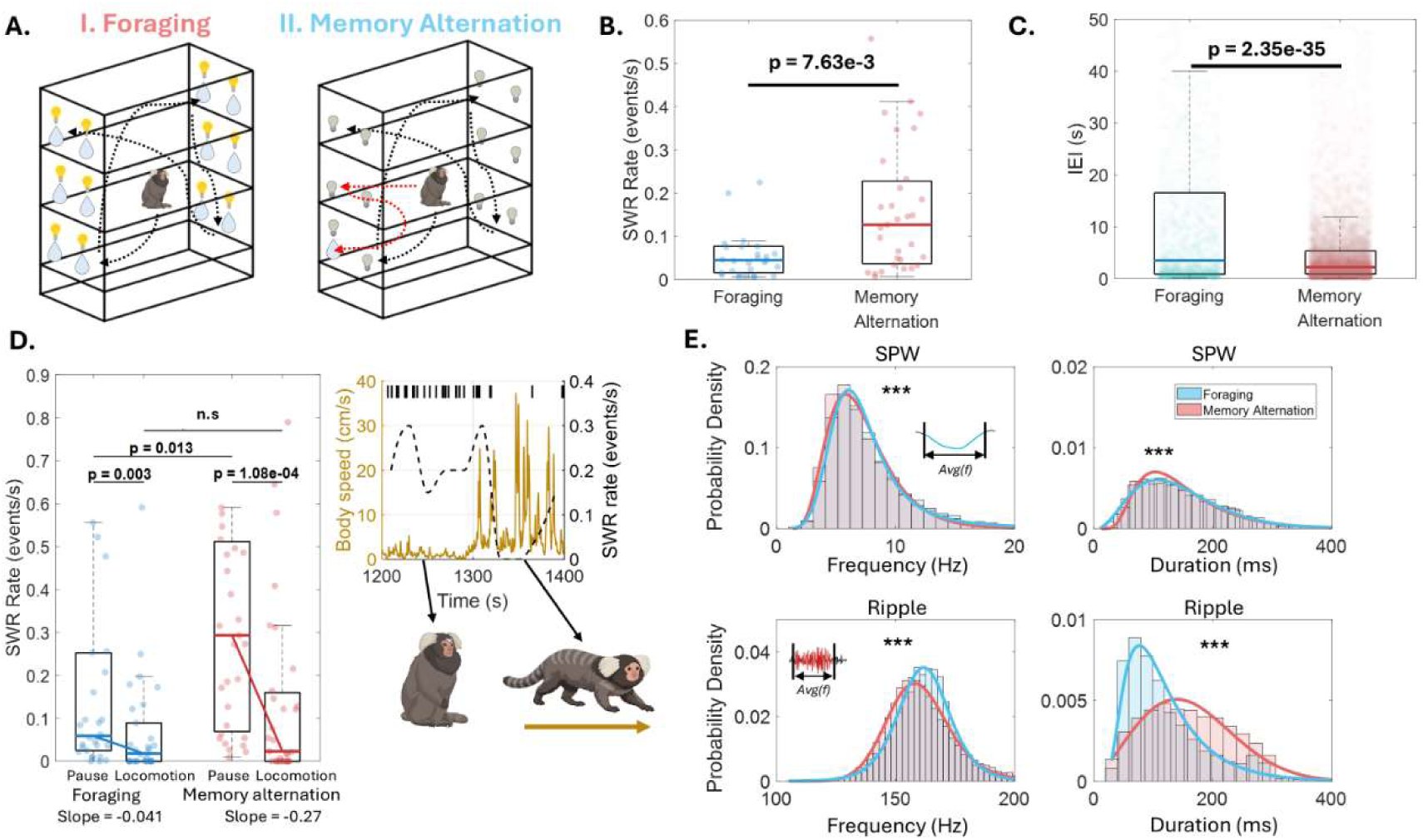
Foraging and memory alternation task differences related to SWR rates, IEIs, body movements and feature distributions. **A.** Illustration of Foraging and memory alternation tasks. Memory Alternation shows the dashed red lines to illustrate the rewarded trajectory for correct reward site sequence. **B.** Session-level SWR rate (events/s) was higher during memory alternation than foraging. **C.** Inter-event intervals (s) were shorter during memory alternation. **D.** SWR rate between pauses and locomotion epochs during the foraging and memory sessions. The task medians are connected by a line, and their slope values are shown beneath the x-labels. Statistics are shown in **Supplementary Table 1.** Left panel shows a recording segment from Subject L (Memory alternation) to showcase the increased SWR rates during pauses versus locomotion. Body speed over time is shown in yellow, with vertical lines on the top representing SWR occurrences and the black dashed line showing the SWR rate (20s window with 5s moving average smoothing). Below are illustrations to depict when the animal is in a pause and then when the animal is locomoting, shown with black arrows. **E.** Distributions of SWR component features (SPW mean frequency and duration; Ripple mean frequency and duration) show task-dependent shifts. Solid lines represent distribution fits to highlight differences in features. Illustrations in the mean-frequency panels emphasize that this measure reflects the average instantaneous frequency of each component. All sample sizes, normality tests (K–S), and rank-sum statistics (tail, p, test statistic) are reported in **Supplementary Table 1**.

During both tasks, SWR rates were higher during pauses than during locomotion (**Fig. 3D** and **Supplementary Table 1**). In foraging sessions, the rate medians were 0.059 SWRs/s versus 0.018 SWRs/s for pauses and locomotion, respectively (Wilcoxon rank-sum, one-tailed, p=0.003), and 0.29 SWRs/s versus 0.02 SWRs/s for pauses and locomotion during the memory alternation sessions (Wilcoxon rank-sum, one-tailed, p=0.0001). **Fig. 3D** (right panel) shows an example of the body and head angular speed traces during periods of locomotion and pauses for subject L alongside SWR occurrence to illustrate this trend.

One potential confounder in these analyses is that animals spent more total time paused than in locomotion. Although the time spent in locomotion was relatively long (4,549.5 s), one may argue that this imbalance could increase the likelihood of observing SWRs during the longer pauses (**Fig. 3D, right panel**). To rule out this confounder, we performed a temporal equalization analysis. For each session, we randomly subsampled body-pause intervals so that the total sampled pause duration matched the total duration of locomotion intervals and repeated this procedure 1,000 times per session to avoid dependence on a single random subsample. After temporal equalization, the duration-matched pause SWR rate remained higher than the locomotion SWR rate. Across sessions, the duration-matched pause SWR rate was 0.2654 events/s on average, compared with 0.1407 events/s during locomotion (Wilcoxon rank-sum test, one-tailed, p = 0.0120). Across the full permutation distribution, the matched pause-minus-locomotion rate difference was 0.1247 events/s, with a 95% permutation CI of [0.0100, 0.3185] and an empirical permutation p-value of 0.012. Thus, the difference in SWR rate was not explained by the longer total duration of pause periods.

It is possible that memory demands not only influenced SWR rates but also SWR features such as duration and frequency. To investigate this issue, we examined whether such features differed between tasks. Small but significant differences were observed in SPW mean frequency (Wilcoxon rank-sum, two-tailed, p = 8.48e-19), SPW duration (Wilcoxon rank-sum, two-tailed, p = 1.82e-4) and Ripple mean frequency (Wilcoxon rank-sum, two-tailed, p =2.5e-22). Notably, Ripple duration was substantially longer during memory alternation than during foraging (Wilcoxon rank-sum, two-tailed, p = 4.8e-282) (**Fig. 3E**). Together, these results indicate that during memory alternation, SWRs occurred more frequently and had longer durations.

### Effect of head movements on SWR rates

A distinctive feature of marmoset behavior is the execution of exploratory rapid head movements that occur more often during pauses (**Fig. 1 and Supplementary Fig. 1**). To investigate how head movements affect SWR rates, we divided the pause periods into two different substates: 1) pause with head stationary, 2) pause with head moving during the two tasks (**Fig. 4A**). SWR rates were significantly higher when the head was stationary (state 1) compared to when the head was moving (state 2), with median rates in foraging sessions of 0.058 versus 0.022 events/s for state 1 and state 2 respectively, and 0.29 versus 0.051 events/s for state 1 and state 2 during the memory alternation sessions. This difference was not significant in foraging sessions (Wilcoxon rank-sum, p = 6.57e-2), but it reached significance during memory alternation sessions (Wilcoxon rank-sum, p = 1.85e-3). Comparing between tasks revealed that state 1 SWR rates were significantly higher in memory than in foraging (Wilcoxon rank-sum, p = 2.53e-2), whereas State 2 rates did not significantly differ between memory and foraging (Wilcoxon rank-sum, p = 1.93e-1). Overall, SWR rates were greater during pauses with head fixations, primarily in memory alternation relative to foraging. This suggests that the enhancement of SWR occurrence during pauses with head fixations is amplified under greater memory demand (**Fig. 4A**).

**Figure 4.**
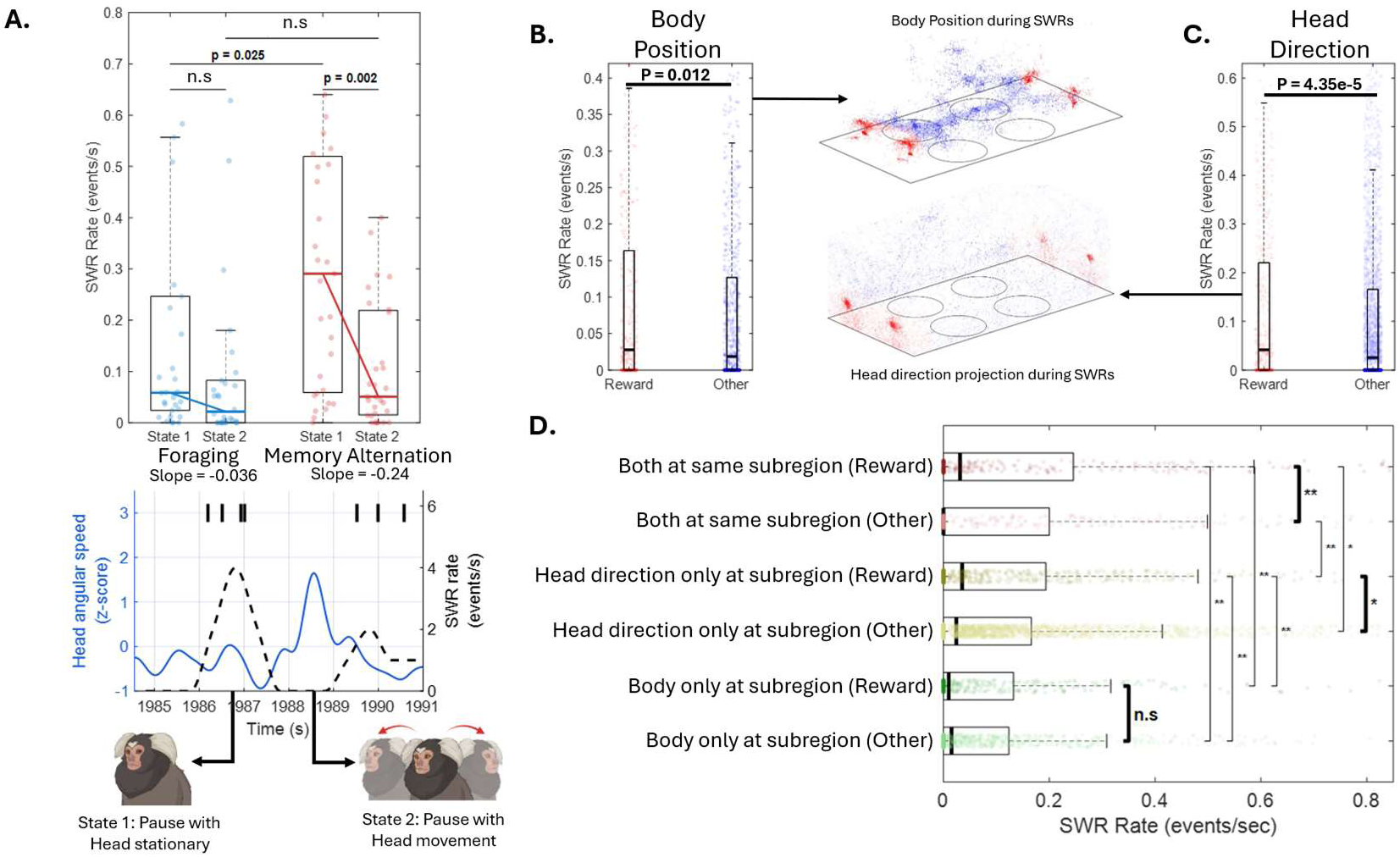
SWRs and their association with head and body movements, as well as their relation to reward subregions. **A.** Boxplots of SWR rates for each of the two states, shown separately for foraging and memory alternation sessions. Median lines are highlighted with thicker task-colored lines and connected to show the slope difference between states (n = 29 foraging sessions, n = 30 memory alternation sessions). Slope values underneath each respective task x-axis label. Bottom panel is a segment of recording from Subject C (Memory alternation task) to showcase the increased SWR rates during head stationary periods compared with when head movement increases. Here head angular speed (z-score) over time is shown in blue, with the ticks on the top representing SWR occurrences and the black dashed line showing the SWR rate (0.5s window with 1s moving average smoothing). Illustration below to demonstrate examples of State 1 and State 2. Pause and head stationary (State 1, left) and pause with head movement (State 2, right) during this segment (See Methods). **B.** The SWR rates for body positions at reward subregions, compared with other subregions (divided into 72 subregions per session; See Methods for maze segmentation and reward versus other classifications) (one-tailed Wilcoxon rank-sum, p = 0.012). **C.** SWR rates when the head direction of the animal pointed toward a subregion with a reward versus other (one-tailed Wilcoxon rank-sum, p = 4.35e-5). In **B** and **C,** the median line of boxplot is highlighted by a thicker black line. To the right of each are panels with the traces of the body position and head direction on the second floor of the maze, red dots marks SWRs occurring at reward subregions and blue marking SWRs occurring in other regions. **D.** To examine the interplay of head direction and body location and the influence this may have on SWR rates, the following conditions were established: 1) Both (head direction and body position) at the same subregion, 2) Head direction only at subregion, and 3) Body only at subregion. We compared these three conditions with the reward versus other subregions (Kruskal-Wallis, p = 3.95e-3, results of the significant post hoc Wilcoxon rank-sum one-tailed test shown in figure axis with asterisk, * p-value < 0.05, ** p-value < 0.01). Median line of boxplot is highlighted by thicker black line. Bold asterisks and lines denote comparisons of interest, showing significant reward effects when both body and head direction were reward-aligned and when only head direction was reward-aligned, but not when only body position was reward-aligned.

### Reward and SWR rates

Previous studies have reported that SWR rates in the HPC of freely navigating rodents increase when a task outcome is associated with reward **^[47]^**. Here, we explore whether the marmoset’s body location relative to reward subregions or their gaze direction (measured by head direction) toward those reward subregions influenced SWR rates. When the animal’s body was positioned at a reward subregion (within an approximate perimeter of 30 cm from reward, equivalent to one body length), SWR rates were significantly elevated compared to other areas (reward mean = 0.129, median = 0.028 vs. other mean = 0.106, median = 0.018; p = 0.012, Wilcoxon rank-sum, **Fig. 4B**). Similarly, when the animal’s head direction was directed at a reward subregion, SWR rates were significantly higher compared to when head direction was directed to non-rewarded subregions (reward mean = 0.148, median = 0.041 vs. other mean = 0.132, median = 0.026; p = 4.35e-5, Wilcoxon rank-sum, **Fig. 4C**).

To further disentangle the contributions of reward-directed head orientation and body position to SWR rates, we grouped SWR rates according to whether the animal’s head direction and/or body position occupied reward-associated subregions. Reward refers to subregions containing reward sites, whereas Other refers to non-reward subregions. We defined six conditions using the labels shown in **Fig. 4D**: Both at same subregion (Reward), when head direction and body position were assigned to the same reward subregion; Both at same subregion (Other), when head direction and body position were assigned to the same non-reward subregion; Head direction only at subregion (Reward) or Head direction only at subregion (Other), when head direction was assigned to a reward or non-reward subregion while body position was assigned to a different subregion; and Body only at subregion (Reward) or Body only at subregion (Other), when body position was assigned to a reward or non-reward subregion while head direction was assigned to a different subregion.

A significant main effect was observed across groups (Kruskal-Wallis test, p = 3.95e-3). Post hoc Wilcoxon rank-sum tests showed that SWR rates were significantly higher for Both at same subregion (Reward) compared with Both at same subregion (Other) (median = 3.12e-2 versus 0.00 events/s, p = 1.43e-3). A similar effect was observed for head direction alone, where Head direction only at subregion (Reward) showed significantly higher SWR rates than Head direction only at subregion (Other) (median = 3.53e-2 versus 2.43e-2 events/s, p = 4.82e-2). In contrast, body position alone did not show a reward-specific increase, as Body only at subregion (Reward) did not differ significantly from Body only at subregion (Other) (median = 9.92e-3 versus 1.53e-2 events/s, p = 7.99e-1). A log-rate GLM using the same rate vectors confirmed this pattern, with reward-associated increases for Both at same subregion (fold change = 1.95, one-tailed p = 3.55e-6) and Head direction only at subregion (fold change = 1.18, one-tailed p = 3.75e-2), but not for Body only at subregion (fold change = 0.87, one-tailed p = 8.51e-1) (See Methods).

Together, these results indicate that SWR rates were highest when both head direction and body position were aligned to the same reward-associated subregion, followed by cases in which head direction alone was aligned with reward. Body position near reward, when head direction was directed elsewhere, did not produce a significant reward-specific increase in SWR rate, suggesting that reward-directed head orientation is a stronger predictor of reward-related SWR modulation than body proximity alone.

## Discussion

We recorded hippocampal SWRs in a freely moving primate, the common marmoset, during two different spatial navigation tasks: 1) foraging and 2) memory alternation. We found that marmosets navigate by alternating between pauses and locomotion, spending considerable time in the former state. During pauses, animals made frequent, rapid head movements to visually explore the maze and reward locations. SWR rates in hippocampal subfields CA1 and CA3 were higher during pauses than during locomotion and higher during the memory task compared to the foraging task. Importantly, during pauses, SWR rates were highest when the animal was near the reward location and its head was oriented toward it.

### A conserved ethological motif for spatial navigation in mammals

In laboratory rodents, spatial navigation is characterized by a recurring behavioral motif in which animals alternate between periods of locomotion associated with active external exploration and brief pauses during which behavior becomes dominated by internal evaluation of space and action. During locomotion, rodents continuously sample the environment through whisking, sniffing, head movements, and visual exploration while moving toward goals, whereas pauses, including stopping, rearing, or hesitation at decision points, are thought to provide opportunities for internal processing, spatial updating, memory retrieval, and route evaluation **^[4,8,48]^**. This alternation between externally driven exploration during movement and internally driven processing during pauses has been proposed as a fundamental ethological motif of rodent behavior during navigation, linking overt movement through space with covert evaluation of cognitive maps and planning of future trajectories **^[24–25,49]^**.

Our observations in freely moving marmosets reveal a somewhat similar motif. Like rodents, marmosets alternate between movement epochs and pauses, suggesting that this motif may represent a conserved mammalian strategy for navigation **^[4,8,48]^**. However, during pauses, marmosets frequently execute rapid head and gaze shifts to foveate relevant regions of the scene **^[11]^**, and they spend substantial amounts of time in these stationary states. This specific behavior suggests that long pauses in primates are active epochs of visually guided, detailed scene evaluation. Primates may require long stationary periods for accurate visual sampling because high-acuity foveal vision is difficult to maintain during locomotion, when body motion and vestibulo-ocular reflexes must continuously stabilize gaze **^[38,50–53]^**. Primates make 3-5 saccades per second during exploration of a scene and fixate for ∼200 milliseconds to obtain a high-resolution but spatially limited sample of the scene’s main elements **^[54]^**. This dynamic may ‘expand’ the time a primate spends during a pause, directing gaze to different parts of the scene. In contrast, rodents can continuously sample the environment during movement through whisking, olfaction, and optic flow across largely afoveate retinas, reducing the need for prolonged stationary visual evaluation **^[55–57]^**. Thus, although the pause-locomotion motif is preserved across species, it is shaped by different sensory sampling strategies, reflecting each species’ distinct ecological niche and evolutionary history **^[28,58–60]^**.

### SWR rates vary during pauses and locomotion

Previous studies in head-restrained macaques have shown that hippocampal SWRs occur during awake visual exploration and are linked to saccadic eye movements **^[61]^**. During visual search tasks, SWRs are aligned with fixation periods, small saccades, and successful target detection **^[62]^**. SWR features also appear to vary with behavioral state **^[35]^**. However, head-restrained studies do not engage the complete primate brain navigational machinery, including vestibular, proprioceptive, and motor systems, making direct comparison with studies in freely moving rodents challenging. A recent study in freely moving marmosets has revealed that neurons in the HPC subfields CA1 and CA3 are tuned to the direction of head (view) and suggested that because vision is dominant in primates, it drives landmark-based navigation **^[11,32,41,63–65]^**and memory formation **^[63]^**. One specific contribution of our study is that we quantified SWRs in a freely moving primate navigating a 3D maze, resembling studies in freely moving rodents **^[5]^**. As in rodent studies **^[5,24,66–67]^**, we found that SWR rates were significantly higher during pauses than during locomotion. These results support the hypothesis that SWR broadcasts information from the HPC to other brain regions during awake states, including periods of locomotion pauses and internally driven exploration **^[12]^**.

Primates use foveations during body stops to explore the environment and orient themselves during spatial navigation tasks **^[11,42,68–69]^**. During such periods of visual sampling, new memories are formed, and old memories of familiar places or objects are retrieved and updated to inform the decision to navigate in a certain direction (e.g., toward a likely-rewarded location). We found that SWR rates were highest during such periods of fixation. One could hypothesize that images of reward locations and landmarks in the 3D maze, acquired during fixations, reactivate memories and trigger SWRs that engage the neocortex-hippocampal memory network **^[14,70–72]^**. Supporting this hypothesis, we found that SWR rates were significantly higher when the animal’s head was directed toward a reward subregion than toward non-rewarded subregions. A similar result has been reported in head-fixed macaques **^[62]^**. Remarkably, SWR rates were elevated when the animal’s body was physically located close to a reward subregion, but mainly when the head was directed to the rewarded locations. The latter suggests that gaze directed toward the rewarded locations is the primary driver of SWR rates.

One possible explanation is that proximity to reward may trigger recall of memories associated with previous rewarded experiences. During navigation, animals must maintain a spatial (cognitive) map of the 3D maze layout in allocentric or egocentric coordinates **^[73]^**, encoding proximity to behaviorally relevant landmarks (i.e., a rewarded location). Interestingly, a study in macaques found that location within a virtual maze and the positions of reward objects modulate the emergence of spatial selectivity in HPC CA3 and CA1 neurons in an egocentric frame **^[32]^**. One study in freely moving humans found that theta oscillations were more prominent when subjects moved and approached the edges of a room; however, their intensity varied with movement speed, and SWRs were not quantified **^[74]^**. Our results suggest that SWRs coupled to active vision (head-direction exploration) may be an essential mechanism for forming and updating cognitive maps during navigation in primates.

### SWR and memory-guided navigation

Contrasting the foraging and memory alternation tasks revealed task-specific differences in SWR dynamics and component features. Session-level SWR rates were higher and inter-event intervals (IEIs) were shorter during memory alternation than during foraging (**Fig. 3**). These results suggest that the higher cognitive demands of memory alternation, requiring internal recall of task structure, space, and recent outcomes, are associated with increased SWR production and tighter event clustering. On the other hand, during foraging, animals may rely more heavily on ongoing sensory exploration and may place comparatively lower demands on internal memory-guided updating. Interestingly, SPW had higher mean frequency (median 6.92 vs 6.56 Hz), and lower duration (median 132 ms vs 135 ms) during foraging than during memory. Ripple mean frequency was slightly lower in memory (median 158.96 vs 162.01 Hz). Most notably, Ripple duration was markedly longer in memory (median 155 vs 100 ms), a difference that could not be explained by the diminished oscillatory frequency of the Ripples. Our results agree with previous studies linking longer Ripple duration to increased memory demands **^[5,15]^**.

### SWR features across species

We independently characterized the low-frequency SPW and high-frequency Ripple components of SWRs in all events as well. Consistent with previous reports in primates, Ripple frequencies typically ranged from 100 to 250 Hz **^[14,33–34,36,43,62,75–77]^**. Both SPW and Ripple duration distributions were positively skewed, with a concentration of events in the 50 – 100 ms range, particularly for the Ripple component. This is similar to observations in human studies, where Ripple durations typically cluster around 50–100 ms, whereas in rodents, durations often span a broader range centered around ∼70 ms.

The SPW component exhibited longer durations and higher power within the examined window than the Ripple component, as expected, given that the SPW typically produces a larger voltage deflection **^[5,78–79]^**. Our Ripple mean frequencies seem higher than those reported in restrained macaques **^[33–34,62]^**, and humans **^[14,80–81]^**, and closer to those reported in rodents **^[5,18–19]^**. These differences may be due to interspecies variability (macaque vs marmoset vs rats) and may be associated with brain size **^[82]^**, or experimental conditions (e.g., freely moving vs. body- head-restrained). It is also possible that our detection algorithm included some high-frequency oscillations (HFO) that coexist with theta waves. This issue is endemic to studies of SWRs **^[4,16,43–44,83–84]^**. More precise detection algorithms linked to controlled behavioral paradigms may be needed to characterize the full spectrum of fast-wave oscillations in primates and their relationship to memory and behavior **^[43–44,85]^**.

Due to variability in detection algorithms and experimental designs, it is difficult to directly compare quantitative values recorded across studies in different species **^[43]^**. However, through validation, we feel that our methodology applies the proper detection constraints proposed by others in the field, ensuring that true SWR events were detected. We demonstrated validation of our detection using real recordings labelled by experts and compared this with three other published methods of SWR detection. The results showed that our method for detecting SWR performs comparably to methods reported in previous studies **^[13,44,46]^**. Additionally, our method can be applied to single-channel data and serve as a foundation for developing AI-based SWR detection methods grounded in marmoset neurophysiological data.

In summary, our study identifies a phylogenetically conserved hippocampal navigation motif that has survived one of the major sensory transitions in mammalian evolution: the shift from predominantly nocturnal ancestors that relied heavily on olfactory and tactile exploration to diurnal primates that navigate through active visual sampling. While the sensory information used to construct spatial representations has changed dramatically, the hippocampal computation associated with transitions between external exploration and internal evaluation appears remarkably conserved. Our findings, therefore, suggest that evolution preserved a fundamental navigation algorithm while reconfiguring its sensory interface to accommodate the visual specializations of primates.

## Materials & Methods

### Animals

Animal care and housing were the same as in Piza et al. **^[11]^**. Three common marmosets (Callithrix jacchus; two males and one female; ages 3, 5, and 8 years) were used. The subjects were housed in customized cages designed for primates and maintained a 12-h light cycle (Night: 7:00 p.m.–7:00 a.m.; Day: 7:00 a.m.–7:00 p.m.). The animals were cared for and maintained within the primate facility at the Robarts Research Institute. The diet of the animals consisted of dry food formula with fruit, nuts, and protein sources to serve as supplementary foods. To be transferred into the plexiglass transfer boxes and navigate the maze a positive reinforcement was administered which consisted of condensed milk or gum Arabic (Acacia) which was either directly administered to the animal or through metal cannulas within the gym. All procedures were under the approval of the Animal Care Committee at Western University (London, Ontario) and in compliance with the Canadian Council for Animal Care (CCAC).

### Experimental Set-up

The experimental set-up follows Piza et al. **^[11]^**and additionally included the memory alternation (sequence) task. Subjects freely navigated a transparent rectangular “gym” with three floors and 12 reward subregions (4 per level; two positioned on opposing sides of each level). Animals were habituated to the apparatus for around 2 weeks. In many sessions, a cage partner was present to reduce stress during free exploration. Task contingencies and reward delivery (solenoid valve) were controlled using NIMH MonkeyLogic **^[86]^**.

### Tasks

1) Foraging task: A single reward site was cued with a flashing LED in a pseudo-random order. A “correct” engagement was registered when the head-tracking marker entered a ∼15 cm radius around the active reward site (the LED cue and reward spout were co-located; for monkey C, the LED was positioned immediately above the sipper tube, and for monkey L the spout itself illuminated). Upon entering this zone, a sound cue was played, and liquid reward (condensed milk) was delivered. Sessions lasted as long as the animal continued to forage (typically around 40–60 min). For subject L, the foraging epoch was shorter on days when the memory task was also run (typically a few minutes). In some PB sessions, hand delivery was used to encourage harvesting, consistent with our earlier training procedures, this animal never learned the LED cue. 2) Memory alternation task: Two reward subregions were selected for that session (i.e., a two-site configuration), and rewards alternated between them such that the subject could not obtain reward at the same site twice in a row without first visiting the other site. During the encoding/guided phase, the currently rewarded target was indicated by a flashing LED.

A trial was defined as a reward-to-reward transition: it began when the animal left the previously rewarded site and ended when it reached the current target site and received reward. Correct responses were therefore defined by selecting the currently rewarded target site (again operationalized as entering within ∼15 cm of that target), which triggered reward delivery and the “correct” sound cue. Incorrect trials correspond to deviations from the intended alternation sequence, including intermediate visits/engagements at non-target locations before arriving at the correct target. Operationally, this was captured as trials that did not satisfy the “direct” target transition (e.g., reaching a non-target region / engaging an off-target spout prior to the correct target), and in some sessions also included a timeout rule (e.g., failure to reach the correct target within a set window). Importantly, in this freely moving paradigm, incorrect trials often reflect variable intermediate deviations rather than immediate trial termination, and trials still typically ended when the animal eventually reached the correct target and received reward.

After performance reached the criterion, the guiding LEDs were turned off, and the animal was required to retrieve rewards based on memory of the two-site configuration and alternation rule. In our analyses and table, “trials to criterion” were computed from the behavioral sequence as the first point at which performance exceeded 60% correct for subject C and 70% correct for subject L, using a 5-trial moving average. **Supplementary Table 2** summarizes all sessions across both tasks (foraging and memory alternation). Across the dataset, 49 recording sessions were analyzed. For task/state analyses, recordings containing both foraging and memory alternation periods were split into separate task-specific epochs, yielding 61 task-specific epochs. For reward-subregion analyses requiring behavioral event codes, the corrupted Session 35 epoch from Subject L was excluded, leaving 59 task-specific epochs.

### Data Acquisition

Data acquisition follows that performed in the Piza et al. 2024 study **^[11]^**. Subjects were implanted with a wireless recording head stage and electrodes covered with a 3D-printed cap. Six spherical retro-reflective markers were placed on the cap using unique geometrical configurations per subject, which allowed for six-degrees of freedom motion capture tracking (Optitrack, Flex 13, Corvallis, OR). Before the session, the setup was calibrated using a calibration wand (CWM-125, Optitrack). This wand uses 3 markers of known dimensions and distance to compute the volumetric coordinate system through a triangulation algorithm. A calibration square (CS-100, Optitrack) was used to define a zero point and the plane corresponding to the ground. The zero-point was always the south-west corner of the maze. This would allow the tracking software (Motive, Optitrack) to compute the position and orientation of the cameras and then subsequently the 3D position of the cap markers to track the subject location (X, Y, Z) as well as the head orientation (roll, pitch, yaw).

For the surgical implantation of the electrodes, co-registration of anatomical MRI (9.4 T) and microCT scans (150 μm) (3D Slicer software, slicer.org) was used to implant the recording chamber. A microbrush array of 32 channels was implanted and the final position of the array was verified with co-registration of post-surgical CT scan to the pre-surgical anatomical MRI. The microwire brush array consisted of platinum-iridium electrodes (Microprobes Inc., MD, USA). The final locations for the electrodes were verified to be in CA1 for subject C and CA3 for subject PB and subject L (**Supplementary Fig. 2**). The individual contact locations cannot be reconstructed but the bundle location and implanted anatomical region could be reconstructed. The raw neural data was sampled at 30 kHz and saved within NS6 files and preprocessed through the Blackrock acquisition system software.

### Signal pre-processing

To extract the LFP signal, the raw signal was downsampled to 1 kHz and a fourth-order zero-phase low-pass filter with a 250 Hz cutoff (filtfilt function, MATLAB) was applied to extract the LFP signal, along with notch filters for power line noise removal at 60, 120, and 180 Hz **^[87]^**. Offline spike sorting was conducted using the methods done in the Piza et al. 2024 study ^[11]^.

### SWR detection

LFPs were recorded using a 32-channel brush array, and SWRs were detected from these signals. Rather than selecting a single “best” channel for analysis, all channels were included after applying standardized noise exclusion criterion. Specifically, channels exhibiting persistent high-frequency noise exceeding 5 standard deviations above baseline were removed from further analyses. Additionally, within each remaining channel, time periods in which the root mean square (RMS) of the signal exceeded 7 standard deviations were excluded to remove transient artifacts, such as movement-related noise. After these exclusion steps, each channel was analyzed independently using the SWR detection pipeline. Following recommendations from Liu et al. (2022) regarding heterogeneity in SWR detection methods **^[43]^**, we combined approaches commonly used for primate SWR detection **^[43,62,75–77]^**. The first step was to take the LFP signal and apply two filters to separate the Ripple and SPW components. For the Ripple component, a 4th order zero phase bandpass filter of 100 to 250 Hz **^[62]^** is applied, while for the SPW, a 1st order zero phase bandpass filter of 1 to 10 Hz is applied. The Ripple amplitude envelope is computed taking the Hilbert transform **^[88]^**of the Ripple component signal post-filtering and taking the absolute squared value. The amplitude envelope is clipped to remove extreme values that are above 5 standard deviations of the amplitude envelope to account for noise. The amplitude envelope is then smoothed using a Kaiser-window finite impulse response low-pass filter with a cutoff frequency of 40 Hz. The SPW signal Hilbert transform was squared and smoothed prior to the threshold computation as well.

The Hilbert transform was computed using MATLAB’s built-in Hilbert function. The threshold was computed by taking the squared and smoothed components and then independently z-scoring both the Ripple envelope and SPW signals across the entire signal. For threshold computations, the Ripple periods of interest were defined as windows where the z-scored Ripple envelope exceeded 3 STD and fell back down to 1 STD, while for the z-scored SPW it was when the STD exceeded 2 and fell back to 1 STD. Windows in which these two conditions occurred within at least 50 ms from one another were classified as a SWR candidate. The coincidence of a SPW and Ripple candidate allowed for higher specificity of classical SWRs rather than Ripple only events **(Fig. 2C)**. One exclusion criterion required Ripple components to last at least 50 ms, like the Leonard and Hoffman 2017 study **^[62]^**, to consider and remove high-gamma noise artifacts. Another exclusion criterion required that the Ripple must contain three peaks in the oscillation above the detection threshold to qualify as a Ripple, like the Norman et al. 2019 study **^[14]^**, to remove possible noise from unit activity that leaks into the LFP signal.

After this automatic detection, a GUI was created within MATLAB to allow for user verification and classification of candidate events into 4 categories: 1) Classical SWR – an event with both a defined Ripple and a SPW deflection, 2) SWR with arbitrary SPW form – an event with a defined Ripple but a SPW that contains an arbitrary form dissimilar to a deflection, 3) Ripple only – defined Ripple with no distinct SPW form, and 4) None – noisy event that contains neither a Ripple nor SPW component. Only user-verified classical SWRs were included in post-processing and feature extraction. We did not apply spectral entropy or correlation-based methods to this dataset because these approaches have primarily been used with linear probes. Due to the fact the exact locations of individual microbrush contacts could not be reconstructed, such spatial methods would be difficult to interpret. **Supplementary Table 3** shows the total event counts across the three marmosets for each of the tasks.

### Feature extraction

Only user-verified classical SWRs were included in post-processing and feature extraction. One of these features was the inter-event time intervals (IEIs), which was calculated as the time between consecutive unique events. As the micro-brush array by design has its electrodes spread more in the transverse plane of where it is injected and the nature of SWRs to travel, it is important to consider the possibility of the 32 channel probes recording the same event on different channels with slight differences in time. Therefore, unique events were defined by collapsing overlapping SWR detections across channels, so that only one event from each overlapping group was retained. This same unique-event definition was used for both SWR rate and IEI calculations, such that SWR rates were computed from the number of unique SWR events per recording duration. For each session, IEIs were calculated between consecutive unique SWR events, using the time from the end of one unique event to the start of the next unique event.

The duration of each component within the event was also extracted. To retrieve this, the start and end indices from the Ripple and SPW are saved during the detection for each event and stored. This allowed for the calculation of the duration of each SWR component by using the difference between the start and end times, along with the sampling rate. The mean frequency of both was also computed using the Hilbert transform of each of the component’s signal in their respective frequency bands from the filtering method mentioned previously (SPW: 1-10 Hz, Ripple: 100-250 Hz). With the Hilbert transform of these signals, we can compute the phase and then the instantaneous frequencies of the component signals. The phase is extracted using the MATLAB angle and unwrap functions, and the instantaneous frequency was computed by taking the difference of the subsequent frequency and solving for:

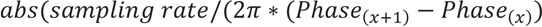

Eq. 1 Instantaneous frequency from the phase of the signal

The instantaneous frequencies were computed for each value within the duration of the SPW and Ripple respectively and then averaged to compute the mean frequency of the components for each of the events.

The spectral components of the events were also analyzed within the frequency and time domain of the LFP signal of the SWRs to understand the interplay of spectral components within the frequency ranges of these events. To achieve this, we utilized MATLAB’s built-in spectrogram function and applied a time window of 400 ms centered at the Ripple peak of the SWR event. We also applied a hamming window of 100 ms as well as signal overlap of 80% to increase the balance in temporal and spectral resolution. The spectrograms were computed for each event, and the power was averaged across all events.

### Parameter fitting

To characterize the distributions of SWR features, we performed parameter fitting for both the mean frequency and duration of the SPW and Ripple components. Frequency distributions were fitted using a stable distribution, while duration distributions were fitted using a gamma distribution. First, the SPW and Ripple mean frequencies were extracted, followed by the computation of their basic statistics, including mean, median, and standard deviation. A stable distribution fit was applied to each frequency dataset using MATLAB’s fitdist function, estimating the four stable parameters: α (stability index), β (skewness), γ (scale), and δ (location). Since most stable distributions do not have a closed-form probability density function (PDF), MATLAB computes the PDF numerically using a direct integration method based on the characteristic equation (Eq.2) **^[89]^**.

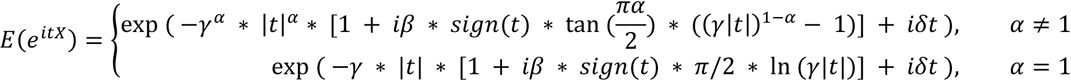

Eq. 2 Characteristic function for the stable distribution

For the duration features, SPW durations were fitted directly using a gamma distribution **^[90]^**, while Ripple durations were first adjusted by subtracting 50 ms before applying the gamma fit. The gamma distribution parameters, a (shape) and b (scale), were estimated using MATLAB’s fitdist function. The fitted gamma PDF was then overlaid onto the normalized histogram of the duration data. To ensure consistency, uniform axis limits were set across plots, and both histograms and fitted distributions were normalized to probability density functions for direct comparison. The resulting fitting parameters were stored for statistical evaluation, and summary statistics were printed to validate the accuracy of the fitted distributions.

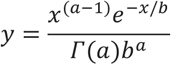

Eq. 3 PDF function for the gamma distribution

### Body and Head movement epoch classifications

Significant head and body movement classifications, and view as facing location were done with the same methods from the Piza et al. 2024 study **^[11]^**. Briefly, Angular head velocity (AHV) was estimated using rotation vectors that were exported from the Motive software as quaternions, and the angular distance (*θ*) between consecutive time points was computed using the MATLAB dist function (Robotics and Autonomous Systems Toolbox). Epochs where AHV exceeded 2000°/s were considered artifacts. Significant head movements were defined as those with an AHV greater than 250°/s and a movement amplitude exceeding 25°. Movement onset and offset were identified as the points where velocity dropped below one-fifth of the peak value. Movements without a clear onset were excluded from further analysis. Locomotion epochs were defined as periods in which translational speed (TS) exceeded 30 cm/s, movement amplitude was at least 30 cm, and duration was at least 1 s. This criterion ensured the analysis of active locomotion involving a change in physical location, rather than transient, in-place movements. Movements with speeds above 300 cm/s were considered artifacts. Head direction was estimated as the facing location, where a ray was cast from the 3D head orientation vector estimated from the trackers on the head stage, and the intersection of this ray with the walls of the maze was defined as head direction. SWRs were categorized into the movement states in **Fig. 3D, and Fig. 4A** based on whether they fell into a significant body or head movement epoch or a pause/stationary epoch that did not meet the significant movement threshold. **Supplementary Table 4** and **Supplementary Table 5** below show the total time spent per session in these movement states as well as a detailed list of the number of epochs and the duration mean, median and std. **Supplementary Table 6** also shows the number of SWRs per state per session along with the computed rate.

### Maze segmentation and SWR rate computation

To compute SWR rates near reward-associated subregions, the maze was segmented into 72 total subregions (Fig. 1A). The North, South, East, and West walls were each divided into 16 subregions, and the Top and Bottom surfaces were each divided into 4 subregions. For body-position analyses, each SWR was assigned to the maze subregion closest to the animal’s tracked X, Y, Z position at the time of the event, based on the Euclidean distance between the animal’s position and the center of each subregion. Reward subregions were defined as the subregions containing reward sites, which corresponded approximately to a spatial scale of one marmoset body length (∼30 cm) around the reward location. Head direction was assigned separately by ray casting from the tracked head-orientation vector to the maze wall surface and identifying the intersected subregion. Each session would have a rate for the 72 subregions. They were separated between the subregion at the reward subregions, and the subregion that did not contain reward subregions. Once this was computed the rates per subregion were compared for the reward versus non-reward subregions independently for body position and head direction (**Fig. 4B,C**).

We also examined the interaction between head direction and body position by comparing cases in which both were assigned to the same subregion, either reward or non-reward, with cases in which only head direction or only body position was assigned to a reward or non-reward subregion while the other variable was assigned to a different subregion (**Fig. 4D**). Occupancy duration for each subregion or condition was computed as the total time during which the animal’s body position, head direction, or combined body/head-direction state was assigned to that bin. Bins with less than 1s occupancy were excluded from rate computation. **Fig. 4A** used 59 task-specific session epochs, whereas **Fig. 4B–D** used 48 recording sessions after one session was excluded due to a corrupted .nev file. For **Fig. 4D**, a complementary GLM was used to verify these results by fitting the log-transformed SWR rates using Eq. 4 below. For each session, subregion, and condition, SWR rate was computed as the number of SWRs divided by the occupancy duration in that condition. Because some session-by-subregion bins contained zero SWRs, rates were log-transformed after adding a small constant, ε, defined as half of the smallest non-zero SWR rate observed across all included bins. ‘Mode’ included Both at same subregion, Head direction only at subregion, and Body only at subregion, and ‘Reward Type’ indicated whether the subregion was Reward or Other. This model tested whether reward-associated subregions predicted increased SWR rates within each mode while accounting for session-level differences.

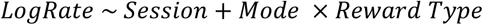

Eq 4. Log transformed GLM equation

### Validation of the Detection Paradigm

Four different SWR-detection methods were tested: the Karlsson detector **^[13]^**, the Kay detector **^[46]^**, Ripple-AI **^[44]^** and our two-threshold detector combining slow-wave (SPW) and Ripple z-score criteria. All detectors were applied to the same continuous LFP, and their detected event onset times were compared against intersecting ground-truth event labels that three independent experts labeled.

For event-level validation, a detected event was considered a true positive if both its onset and offset fell within ±25 ms of the corresponding ground-truth event onset and offset. Ground-truth SWRs that were not matched by a detected event were counted as false negatives. To compute false-positive rate and specificity for ROC analysis, detector outputs and expert labels were also converted into binary time-sample vectors. Time samples outside expert-labeled SWR intervals that were marked as SWR by a detector were counted as false positives, whereas unlabeled samples correctly left unmarked were counted as true negatives. Sensitivity, specificity, and false-positive rate (FPR) were then computed across detection thresholds, and receiver-operating-characteristic (ROC) curves were generated.

The Kay, Karlsson and Ripple-AI detectors were evaluated across Ripple z-score thresholds ranging from 1.0 to 10.0. Our detector was evaluated across a two-dimensional grid of SPW and Ripple z-score thresholds, SPW ranging from 0 to 5, and Ripple ranging from 1.0 to 10.0. ROC curves were constructed using pairs (TPR, FPR), and the area under the curve (AUC) was computed using trapezoidal integration over the ROC points. This provided a complete operating-point comparison between the classical detectors and our SPW and Ripple coincidence approach.

## Resource availability

### Data and Code Availability

The raw and processed data are freely available at this directory: (https://web.eng.fiu.edu/jrieradi/SWR_Code_NMDLab/). Along with the raw files is the MATLAB based code used for the SWR detection, as well as post-processing scripts. The folder also contains the dependencies to run the detection and post-processing scripts such as the Blackrock Neurotech neural processing MATLAB kit **^[91]^**, Marmoset 3D gym scripts **^[11]^**, and EEGLab **^[92]^**. The entire folder download is around 266 GB. Read the ‘Readme’ file for further instructions to get started with the detection and analysis scripts.

## Acknowledgements

We would like to thank Dr. Felix Carbonell and Dr. Wensong Wu for their assistance with questions regarding which statistical applications to use. We thank registered veterinary technicians Kim Thomaes and Kristy Gibbs from Western University for technical assistance in surgeries and animal care; Kevin Barker from Neuronitek for design and manufacturing of implantable recording chambers and experimental setup; Jonathan C. Lau from the Division of Neurosurgery at Western University for providing advice in neuro-navigation and targeting of deep brain structures; and Stephen Frey from Rogue Research, Montreal, Canada for providing technical advice regarding microelectrode array implantation, neuro-navigation and manufacturing of custom microdrives. We also thank Emily Otero for proofreading the manuscript. Research reported in this publication was supported by the Canadian Institute of Health Research Project Grant (CIHR); Natural Sciences and Engineering Research Council of Canada (NSERC); Provincial Endowed Academic Chair in Autism; Canada Foundation for Innovation (CFI); Western University BrainsCAN award grant and Healthy Brains, Healthy Lives (HBHL); National Institute of Environmental Health Sciences (NIEHS) under award number T32ESO33955; Wallace Coulter Foundation.

