## Supplementary tables and figures for "A Conserved Mammalian Hippocampal Navigation Motif Reconfigured for Primate Vision"

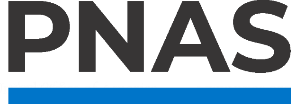

**Supporting Information for**

A Conserved Mammalian Hippocampal Navigation Motif Reconfigured for Primate Vision

*Carlos Otero^1^, Diego B. Piza^2^, Ehsan Aboutorabi^2^, Julio C. Martinez-Trujillo^2^, Jorge Riera^1^

^1^ Biomedical Engineering Dept, Florida International Univ., Miami, FL; ^2^ Schulich School of Medicine and Dentistry, Department of Physiology and Pharmacology, Psychiatry and Neurological Sciences, and Western Institute for Neuroscience, Western University, London, ON, Canada

Corresponding Author: Jorge Riera

**This PDF file includes:**

Figures S1 to S3

Tables S1 to S6

Figures

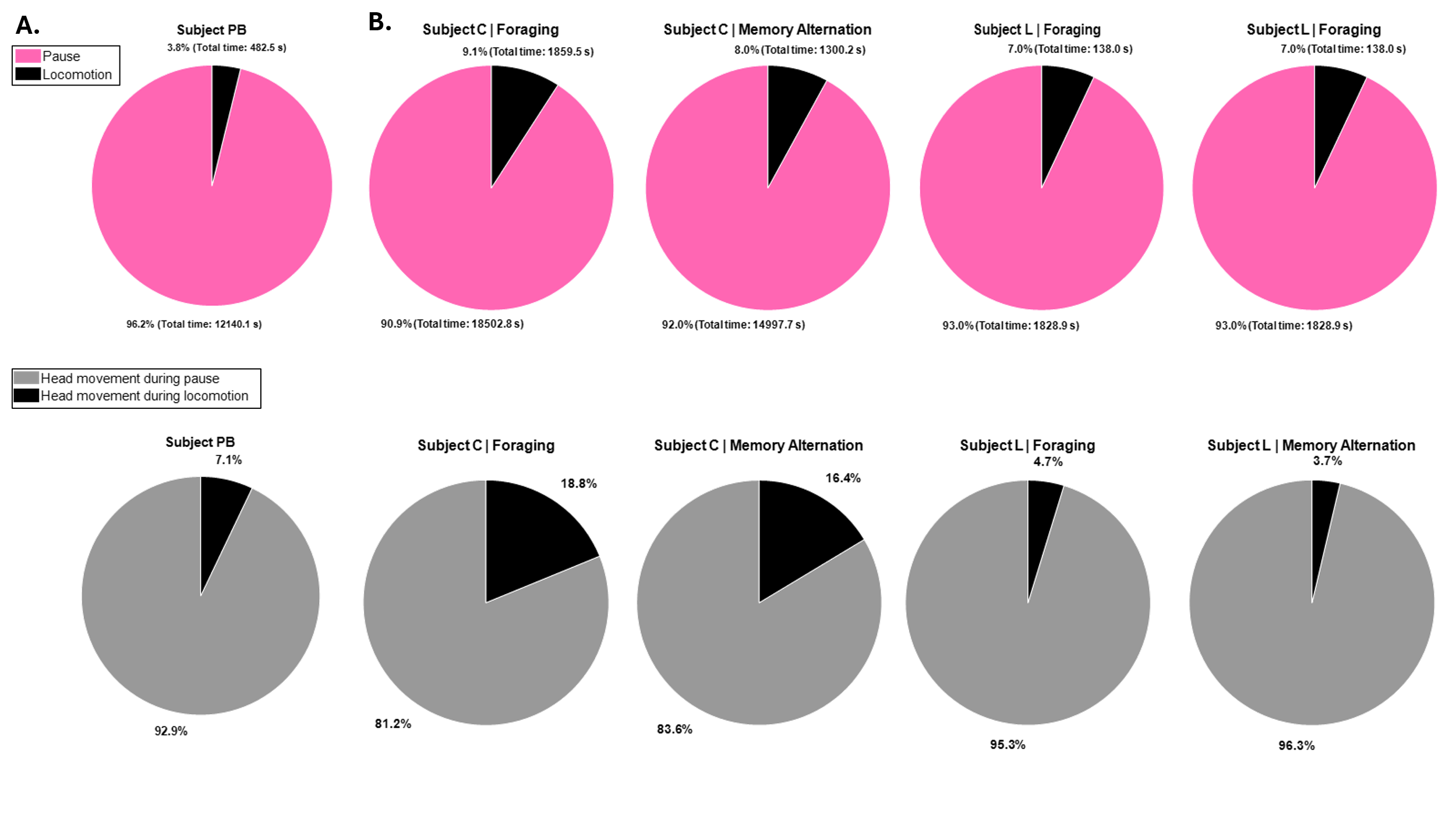

**Supplementary Figure 1.** Pie graphs that depict the proportion of time spent in movement states. **A.** The proportion of time spent in locomotion and pauses across Subject PB (animal who performed only foraging task) sessions (top) as well as proportion of head movements during pauses and locomotion (bottom). **B.** Same as A, but shown separately by task for subjects who performed both tasks (Subject C and L).

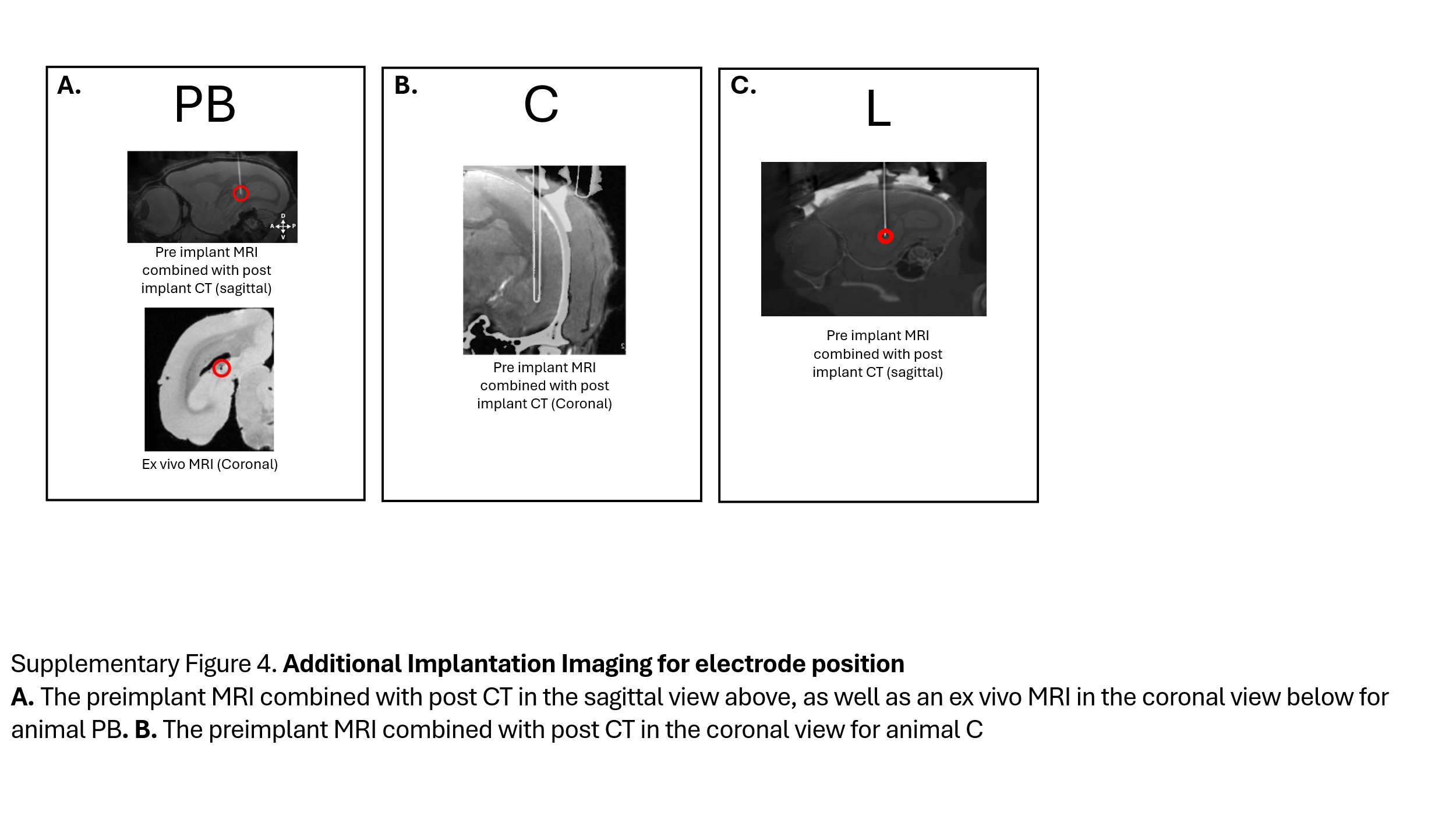

**Supplementary Figure 2.** Additional Implantation Imaging for electrode position in the three marmosets. **A.** The preimplant MRI combined with post CT in the sagittal view above, as well as an ex vivo MRI in the coronal view below for animal PB. **B.** The preimplant MRI combined with post CT in the coronal view for animal C. **C.** The preimplant MRI combined with post CT in the sagittal view for animal L.

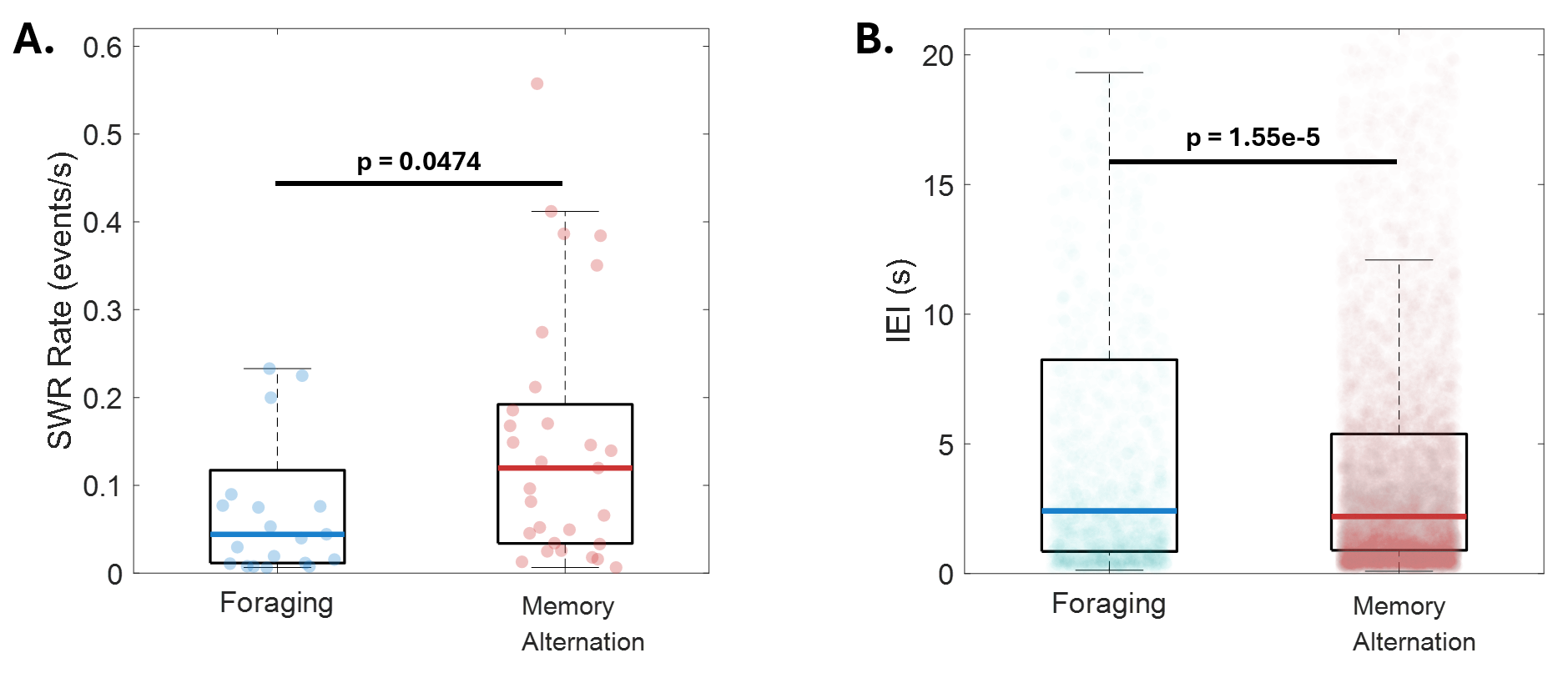

Supplementary Figure 3. SWR rate and IEI comparisons for only subjects that performed both tasks (C and L). A. SWR rates for foraging sessions (n = 21, mean = 0.12 events/s, median = 0.04 events/s) compared to memory alternation sessions (n = 29, mean = 0.15 events/s, median = 0.12 events/s). Memory alternation sessions showed a statistically greater SWR rate median (Wilcoxen rank-sum, one-tailed, p = 0.0474) B. Inter-event intervals for foraging sessions (n = 1881, mean = 16.66 s, median = 2.42 s) compared to memory alternation sessions (n = 9902, mean = 6.91 s, median = 2.20 s). Memory alternation sessions showed significantly shorter IEIs than foraging sessions (Wilcoxon rank-sum, one-tailed, p = 1.55e-5).

Tables

| **Figure** | **N1 (F)** | **N2 (M)** | **K-S**  **P-value**  **(N1)** | **K-S**  **test statistic (N1)** | **K-S**  **P-value (N2)** | **K-S**  **test statistic (N2)** | **Statistical Test** | **tail** | **P-value** | **Test statistic** |
| --- | --- | --- | --- | --- | --- | --- | --- | --- | --- | --- |
| **3B) SWR Rate (events/s)** | 29 | 30 | 5.01e-4 | 0.36783 | 0.411 | 0.15416 | Wilcoxon rank-sum | one-tailed  (M > F) | 7.63e-3 | -2.426 |
| **3C) IEI (s)** | 2291 | 10282 | 1.39e-256 | 0.3581 | <1E-300 | 0.38307 | Wilcoxon rank-sum | one-tailed  (M < F) | 2.35e-32 | 11.785 |
| **3D) SWR rate during locomotion versus pause: Foraging only** | 29 | N/A | 8.95E-04 (pause) | 0.349196 | 9.01E-03  (locomotion) | 0.292689 | Wilcoxon rank-sum | one-tailed (pause > locomotion) | 3.00e-03 | 2.748 |
| **3D) SWR rate during locomotion versus pause : Memory only** | N/A | 30 | 2.58E-02  (pause) | 0.258431 | 2.57E-02  (locomotion) | 0.258581 | Wilcoxon rank-sum | one-tailed (pause > locomotion) | 1.08e-04 | 3.698846 |
| **3D) SWR rate during locomotion versus pause : Pause, Foraging vs Memory** | 29 | 30 | 8.95E-04 | 0.3492 | 2.58E-02 | 0.25843 | Wilcoxon rank-sum | two-tailed | 1.28e-02 | -2.4886 |
| **3D) SWR rate during locomotion versus pause : Locomotion, Foraging vs Memory** | 29 | 30 | 9.01E-03 | 0.29269 | 2.57E-02 | 0.25858 | Wilcoxon rank-sum | two-tailed | 2.91e-01 | -1.0569 |
| **3E) SPW mean frequency (Hz)** | 6354 | 13127 | 2.2e-74 | 0.12433 | 2.23e-116 | 0.10156 | Wilcoxon rank-sum | two-tailed | 8.48e-19 | 8.8536 |
| **3E) SPW duration (ms)** | 6354 | 13127 | 1.34e-28 | 0.076739 | 1.11e-69 | 0.078488 | Wilcoxon rank-sum | two-tailed | 0.000182 | -3.743 |
| **3E) Ripple mean frequency (Hz)** | 6354 | 13127 | 4.74e-09 | 0.042459 | 4.72e-35 | 0.055499 | Wilcoxon rank-sum | two-tailed | 2.5e-22 | 9.719 |
| **3E) Ripple duration (ms)** | 6354 | 13127 | 2.72e-88 | 0.13552 | 2.26e-27 | 0.048959 | Wilcoxon rank-sum | two-tailed | 4.8e-282 | -35.888 |

**Supplementary Table 1.** Statistical testing for Fig. 3. This table summarizes the statistical workflow used for **Fig. 3 B–E** metrics comparing Foraging (indicated by “F” in the table) vs Memory alternation (indicated by “M” in the table). For each metric (row), the table reports the sample sizes (n foraging, n memory), Kolmogorov–Smirnov (K–S) normality test results for each group (p-value and test statistic), and the final statistical test used (e.g., t-test or Wilcoxon rank-sum), including the test tail (one- or two-tailed), p-value, and test statistic.

| **Session number** | **Session name** | **Subject** | **Task performed in session** | **Total detected SWRs after manual curation** | **Sequence (reward sites)** | **Foraging start time (s)** | **Memory start time (s)** | **Total Trials conducted** | **Trials to reach criterion** | **Foraging performance (%)** | **Memory performance (%)** | **#correct trials foraging** | **#miss trials foraging** | **#correct trials memory** | **#miss trials memory** | **Notes** |
| --- | --- | --- | --- | --- | --- | --- | --- | --- | --- | --- | --- | --- | --- | --- | --- | --- |
| 1 | 20190222 | C | F | 54 | N/A | N/A | N/A | 13 | 1 | 92.3077 | N/A | 12 | 1 | N/A | N/A |  |
| 2 | 20190225 | C | F | 217 | N/A | N/A | N/A | 70 | 1 | 95.7143 | N/A | 67 | 3 | N/A | N/A |  |
| 3 | 20190226 | C | F | 160 | N/A | N/A | N/A | 68 | 1 | 98.5294 | N/A | 67 | 1 | N/A | N/A |  |
| 4 | 20190227 | C | F | 135 | N/A | N/A | N/A | 59 | 1 | 88.1356 | N/A | 52 | 7 | N/A | N/A |  |
| 5 | 20190228 | C | F | 42 | N/A | N/A | N/A | 74 | 1 | 85.1351 | N/A | 63 | 11 | N/A | N/A |  |
| 6 | 20190301 | C | F | 33 | N/A | N/A | N/A | 56 | 4 | 96.4286 | N/A | 54 | 2 | N/A | N/A |  |
| 7 | 20190305 | C | F | 100 | N/A | N/A | N/A | 118 | 1 | 85.5932 | N/A | 101 | 17 | N/A | N/A |  |
| 8 | 20200925 | C | F | 29 | N/A | N/A | N/A | 151 | 31 | 22.5166 | N/A | 34 | 117 | N/A | N/A | Sites were placed on opposite sides likely cause of low performance |
| 9 | 20201009 | C | F | 39 | N/A | N/A | N/A | 147 | 92 | 29.932 | N/A | 44 | 103 | N/A | N/A | Sites were placed on opposite sides likely cause of low performance |
| 10 | 20210126 | PB | F | 187 | N/A | N/A | N/A | N/A | N/A | N/A | N/A | N/A | N/A | N/A | N/A | Hand delivery so no .nev file |
| 11 | 20190429 | C | M | 730 | [2,4] | N/A | N/A | 64 | 2 | N/A | 94.26 | N/A | N/A | 82 | 5 |  |
| 12 | 20190521 | C | M | 557 | [1,2] | N/A | N/A | 75 | 2 | N/A | 93.3 | N/A | N/A | 70 | 5 |  |
| 13 | 20190527 | C | M | 73 | [3,1] | N/A | N/A | 92 | 1 | N/A | 97.83 | N/A | N/A | 90 | 2 |  |
| 14 | 20190530 | C | M | 31 | [1,4] | N/A | N/A | 31 | 1 | N/A | 87.1 | N/A | N/A | 27 | 4 |  |
| 15 | 20190603 | C | M | 602 | [1,3] | N/A | N/A | 109 | 6 | N/A | 92.66 | N/A | N/A | 101 | 8 |  |
| 16 | 20190604 | C | M | 158 | [3,4] | N/A | N/A | 146 | 9 | N/A | 94.52 | N/A | N/A | 138 | 8 |  |
| 17 | 20201228 | PB | F | 331 | N/A | N/A | N/A | N/A | N/A | N/A | N/A | N/A | N/A | N/A | N/A | Hand delivery so no .nev file |
| 18 | 20210109 | PB | F | 769 | N/A | N/A | N/A | N/A | N/A | N/A | N/A | N/A | N/A | N/A | N/A | Hand delivery so no .nev file |
| 19 | 20210110 | PB | F | 576 | N/A | N/A | N/A | N/A | N/A | N/A | N/A | N/A | N/A | N/A | N/A | Hand delivery so no .nev file |
| 20 | 20210111 | PB | F | 901 | N/A | N/A | N/A | N/A | N/A | N/A | N/A | N/A | N/A | N/A | N/A | Hand delivery so no .nev file |
| 21 | 20210118 | PB | F | 39 | N/A | N/A | N/A | N/A | N/A | N/A | N/A | N/A | N/A | N/A | N/A | Hand delivery so no .nev file |
| 22 | 20210103 | PB | F | 351 | N/A | N/A | N/A | N/A | N/A | N/A | N/A | N/A | N/A | N/A | N/A | Hand delivery so no .nev file |
| 23 | 20210105 | PB | F | 460 | N/A | N/A | N/A | N/A | N/A | N/A | N/A | N/A | N/A | N/A | N/A | Hand delivery so no .nev file |
| 24 | 20210119 | PB | F | 198 | N/A | N/A | N/A | N/A | N/A | N/A | N/A | N/A | N/A | N/A | N/A | Hand delivery so no .nev file |
| 25 | 250606 | L | F & M | 153 (F=0, M=153) | [3,9] | 114.549 | 193.345 | 30 | 8 | 88 | 59 | 7 | 1 | 13 | 9 |  |
| 26 | 250609 | L | F & M | 61 (F=3, M=58) | [10,4] | 15.5229 | 90.6618 | 32 | 7 | 100 | 80 | 7 | 0 | 20 | 5 |  |
| 27 | 250610 | L | M | 536 (F=0, M=536) | [10,4] | N/A | 16.187 | 48 | N/A | N/A | 77 | N/A | N/A | 37 | 11 |  |
| 28 | 250611 | L | M | 306 (F=0, M=306) | [10,4] | N/A | 20.5023 | 41 | N/A | N/A | 70 | N/A | N/A | 29 | 12 |  |
| 29 | 250616 | L | F & M | 1085 (F=97, M=988) | [3,10] | 14.8334 | 85.4263 | 41 | 7 | 100 | 90 | 7 | 0 | 31 | 3 |  |
| 30 | 250617 | L | M | 784 (F=0, M=784) | [3,10] | N/A | 38.9843 | 42 | N/A | N/A | 86 | N/A | N/A | 36 | 6 |  |
| 31 | 250619 | L | F & M | 170 (F=8, M=162) | [3,9] | 20.8228 | 124.649 | 33 | 7 | 100 | 73 | 7 | 0 | 19 | 7 |  |
| 32 | 250620 | L | M | 150 (F=0, M=150) | [3,9] | N/A | 15.0747 | 34 | N/A | N/A | 76 | N/A | N/A | 26 | 8 |  |
| 33 | 250623 | L | F & M | 113 (F=1, M=112) | [4,9] | 14.5185 | 139.658 | 55 | 11 | 64 | 80 | 7 | 4 | 35 | 9 |  |
| 34 | 250624 | L | M | 1042 (F=0, M=1042) | [4,9] | N/A | 25.161 | 39 | N/A | N/A | 77 | N/A | N/A | 30 | 9 |  |
| 35 | 250626 | L | F & M | 1830 (F=1255, M=575) | [3,4] | 29.1471 | 439.285 | N/A | N/A | N/A | N/A | N/A | N/A | N/A | N/A | corrupt .nev file. so behavioral codes are not recorded. |
| 36 | 250708 | L | F & M | 1137 (F=185, M=952) | [3,4] | 98.2657 | 275.162 | 41 | 10 | 70 | 77 | 7 | 3 | 24 | 7 |  |
| 37 | 250709 | L | M | 410 (F=0, M=410) | [3,4] | N/A | 88.1677 | 30 | N/A | N/A | 87 | N/A | N/A | 26 | 4 |  |
| 38 | 250710 | L | F & M | 258 (F=23, M=235) | [3,10] | 360.6554 | 1014.49 | 37 | 23 | 60 | 70 | 11 | 12 | 9 | 5 |  |
| 39 | 250711 | L | M | 593 (F=0, M=593) | [3,10] | N/A | 14.0619 | 46 | N/A | N/A | 89 | N/A | N/A | 41 | 5 |  |
| 40 | 250714 | L | F & M | 256 (F=24, M=232) | [10,4] | 34.6156 | 314.753 | 36 | 9 | 78 | 92 | 7 | 2 | 25 | 2 |  |
| 41 | 250715 | L | M | 426 (F=0, M=426) | [10,4] | N/A | 70.5166 | 48 | N/A | N/A | 85 | N/A | N/A | 41 | 7 |  |
| 42 | 250717 | L | M | 123 (F=0, M=123) | [4,9] | N/A | 59.0325 | 33 | N/A | N/A | 94 | N/A | N/A | 31 | 2 |  |
| 43 | 250719 | L | M | 48 (F=0, M=48) | [3,9] | N/A | 37.6297 | 36 | N/A | N/A | 100 | N/A | N/A | 36 | 0 |  |
| 44 | 250724 | L | F & M | 320 (F=105, M=215) | [3,9] | 16.5801 | 516.895 | 16 | 13 | 55 | 100 | 7 | 6 | 3 | 0 |  |
| 45 | 250725 | L | M | 952 (F=0, M=952) | [10,9] | N/A | 18.042 | 24 | N/A | N/A | 92 | N/A | N/A | 22 | 2 |  |
| 46 | 250728 | L | F & M | 93 (F=7, M=86) | [10,9] | 33.919 | 100.734 | 42 | 8 | 87 | 97 | 7 | 1 | 33 | 1 |  |
| 47 | 250729 | L | M | 50 (F=0, M=50) | [4,9] | N/A | 51.132 | 31 | N/A | N/A | 100 | N/A | N/A | 31 | 0 |  |
| 48 | 250730 | L | F & M | 717 (F=25, M=692) | [4,9] | 18.2199 | 129.345 | 57 | 10 | 70 | 91 | 7 | 3 | 43 | 4 |  |
| 49 | 250731 | L | M | 1096 (F=0, M=1096) | [3,9] | N/A | 17.303 | 29 | N/A | N/A | 93 | N/A | N/A | 27 | 2 |  |

**Supplementary Table 2**. This table summarizes 49 recording sessions (Subjects C, PB, and L) with behavioral performance metrics and total SWR counts after manual curation. Columns include: session identifiers (Session #, date/session name), subject, task, SWR count, sequence/reward-site configuration (when applicable), task start times (Foraging start, Memory start, when applicable), trial totals (total trials, trials-to-criterion), performance (% correct) and outcome counts (# correct / # miss) for foraging and memory, plus notes. F = Foraging and M = Memory alternation. Several Subject L sessions include both tasks in the same recording (F & M), with SWR totals reported as split counts. Entries marked N/A indicate data not collected or not applicable for that session/task

| Subject | Area of Implantation | Foraging SWRs (sessions) | Memory Alternation SWRs  (sessions) | Total SWRs detected |
| --- | --- | --- | --- | --- |
| PB | Right HPC  CA3 | 3812  (9 sessions) | N/A | 3812 |
| C | Left HPC  CA1 | 809  (9 sessions) | 2151  (6 sessions) | 2960 |
| L | Right HPC  CA3 | 1733  (12 sessions) | 10976  (25 sessions) | 12709 |

**Supplementary Table 3**. Overall SWR event totals by subject and task. Manually curated SWR counts (with number of sessions in parentheses) for foraging and memory alternation, along with implant hemisphere/subfield and the combined total per subject.

| **Session Number** | **Monkey** | **Task** | **Total Time Locomotion(s)** | **Total Time Pauses(s)** | **Locomotion Epochs** | **Locomotion Median Epoch Time (s)** | **Locomotion Mean Epoch Time (s)** | **Locomotion STD Epoch Time (s)** | **Pause Epochs** | **Pause Median Epoch Time (s)** | **Pause Mean Epoch Time (s)** | **Pause STD Epoch Time (s)** |
| --- | --- | --- | --- | --- | --- | --- | --- | --- | --- | --- | --- | --- |
| 1 | C | F | 39.4 | 1542.2 | 49 | 0.750 | 0.804 | 0.276 | 69 | 3.450 | 22.350 | 118.932 |
| 2 | C | F | 342.2 | 2651.7 | 387 | 0.750 | 0.884 | 0.359 | 498 | 3.117 | 5.325 | 7.752 |
| 3 | C | F | 218.3 | 2669.9 | 258 | 0.750 | 0.846 | 0.330 | 339 | 3.366 | 7.876 | 19.937 |
| 4 | C | F | 119.6 | 2086.5 | 141 | 0.750 | 0.848 | 0.344 | 219 | 4.600 | 9.528 | 20.894 |
| 5 | C | F | 124.6 | 1883.9 | 147 | 0.750 | 0.847 | 0.321 | 227 | 4.416 | 8.299 | 16.147 |
| 6 | C | F | 154.5 | 356.0 | 167 | 0.800 | 0.925 | 0.360 | 2 | 178.000 | 178.000 | 25.456 |
| 7 | C | F | 354.2 | 2415.6 | 365 | 0.800 | 0.970 | 0.469 | 472 | 2.683 | 5.118 | 12.346 |
| 8 | C | F | 337.4 | 2311.3 | 349 | 0.833 | 0.967 | 0.438 | 484 | 2.766 | 4.775 | 6.534 |
| 9 | C | F | 169.4 | 2585.6 | 160 | 0.958 | 1.058 | 0.476 | 252 | 4.583 | 10.260 | 28.807 |
| 10 | PB | F | 26.2 | 2011.9 | 24 | 0.900 | 1.094 | 0.544 | 47 | 13.916 | 42.806 | 115.936 |
| 11 | C | M | 153.1 | 1095.7 | 159 | 0.817 | 0.963 | 0.428 | 240 | 2.067 | 4.565 | 6.968 |
| 12 | C | M | 130.7 | 2007.8 | 147 | 0.733 | 0.889 | 0.388 | 219 | 4.833 | 9.168 | 12.640 |
| 13 | C | M | 281.7 | 2526.0 | 281 | 0.867 | 1.002 | 0.497 | 374 | 3.625 | 6.754 | 10.890 |
| 14 | C | M | 93.3 | 2300.1 | 104 | 0.758 | 0.897 | 0.405 | 158 | 9.066 | 14.558 | 18.983 |
| 15 | C | M | 336.0 | 2990.7 | 282 | 0.950 | 1.191 | 0.631 | 344 | 4.991 | 8.694 | 16.038 |
| 16 | C | M | 305.4 | 4077.6 | 305 | 0.933 | 1.001 | 0.393 | 390 | 5.416 | 10.455 | 20.533 |
| 17 | PB | F | 213.1 | 1914.3 | 215 | 0.900 | 0.991 | 0.404 | 339 | 2.650 | 5.647 | 10.756 |
| 18 | PB | F | 50.0 | 1586.3 | 45 | 1.050 | 1.110 | 0.475 | 97 | 11.783 | 16.354 | 16.901 |
| 19 | PB | F | 32.1 | 1028.1 | 41 | 0.717 | 0.784 | 0.259 | 69 | 6.583 | 14.900 | 35.262 |
| 20 | PB | F | 5.4 | 430.7 | 6 | 0.908 | 0.905 | 0.350 | 19 | 17.815 | 22.668 | 22.169 |
| 21 | PB | F | 38.0 | 1412.2 | 43 | 0.767 | 0.884 | 0.436 | 86 | 9.283 | 16.421 | 20.774 |
| 22 | PB | F | 64.2 | 1559.2 | 70 | 0.817 | 0.917 | 0.356 | 141 | 8.383 | 11.058 | 12.341 |
| 23 | PB | F | 9.0 | 1033.3 | 10 | 0.892 | 0.902 | 0.340 | 31 | 21.399 | 33.332 | 33.147 |
| 24 | PB | F | 44.4 | 1164.1 | 45 | 0.917 | 0.988 | 0.393 | 74 | 8.933 | 15.731 | 23.336 |
| 25 | L | F | 13.2 | 68.1 | 7 | 1.967 | 1.890 | 0.584 | 10 | 6.050 | 6.811 | 3.990 |
| 26 | L | M | 12.3 | 321.7 | 12 | 0.958 | 1.026 | 0.407 | 14 | 12.949 | 22.978 | 38.515 |
| 27 | L | F | 4.6 | 66.8 | 6 | 0.725 | 0.761 | 0.215 | 6 | 12.426 | 11.130 | 6.364 |
| 28 | L | M | 10.4 | 1103.6 | 10 | 0.942 | 1.045 | 0.527 | 11 | 19.699 | 100.331 | 183.517 |
| 29 | L | M | 43.2 | 1119.0 | 46 | 0.842 | 0.940 | 0.406 | 58 | 10.708 | 19.294 | 25.905 |
| 30 | L | M | 66.3 | 1605.6 | 64 | 0.783 | 1.035 | 0.495 | 76 | 7.841 | 21.127 | 45.869 |
| 31 | L | F | 5.8 | 67.1 | 5 | 1.250 | 1.160 | 0.580 | 6 | 9.458 | 11.191 | 7.402 |
| 32 | L | M | 20.2 | 1802.9 | 24 | 0.700 | 0.840 | 0.341 | 22 | 34.322 | 81.948 | 142.990 |
| 33 | L | M | 7.1 | 1818.6 | 9 | 0.633 | 0.787 | 0.337 | 19 | 24.532 | 95.717 | 205.202 |
| 34 | L | F | 8.6 | 96.2 | 9 | 0.717 | 0.957 | 0.394 | 11 | 7.549 | 8.742 | 6.756 |
| 35 | L | M | 10.5 | 1295.8 | 14 | 0.608 | 0.748 | 0.243 | 14 | 24.515 | 92.560 | 166.820 |
| 36 | L | M | 9.8 | 833.4 | 9 | 0.983 | 1.091 | 0.589 | 12 | 21.273 | 69.452 | 99.532 |
| 37 | L | F | 23.4 | 100.5 | 14 | 1.775 | 1.670 | 0.462 | 16 | 5.933 | 6.280 | 3.072 |
| 38 | L | M | 115.2 | 2081.2 | 85 | 1.367 | 1.355 | 0.582 | 129 | 6.183 | 16.134 | 61.607 |
| 39 | L | M | 15.8 | 1840.0 | 15 | 1.083 | 1.054 | 0.416 | 19 | 17.165 | 96.843 | 216.777 |
| 40 | L | F | 5.1 | 311.3 | 5 | 0.700 | 1.013 | 0.480 | 2 | 155.642 | 155.642 | 149.401 |
| 41 | L | M | 0.7 | 459.3 | 1 | 0.667 | 0.667 | 0.000 | 1 | 459.279 | 459.279 | 0.000 |
| 42 | L | M | 2.4 | 431.1 | 4 | 0.583 | 0.612 | 0.098 | 5 | 25.898 | 86.213 | 117.041 |
| 43 | L | F | 4.6 | 280.4 | 5 | 0.933 | 0.913 | 0.358 | 4 | 78.744 | 70.096 | 54.841 |
| 44 | L | M | 5.0 | 822.4 | 6 | 0.767 | 0.839 | 0.342 | 8 | 74.161 | 102.798 | 84.966 |
| 45 | L | M | 70.2 | 1914.9 | 53 | 1.317 | 1.325 | 0.451 | 58 | 11.549 | 33.015 | 116.243 |
| 46 | L | F | 30.0 | 261.7 | 28 | 0.975 | 1.073 | 0.427 | 25 | 7.383 | 10.468 | 12.899 |
| 47 | L | M | 18.4 | 174.2 | 21 | 0.650 | 0.878 | 0.432 | 11 | 13.366 | 15.837 | 13.353 |
| 48 | L | M | 80.2 | 1580.6 | 62 | 1.300 | 1.293 | 0.484 | 74 | 9.649 | 21.359 | 46.574 |
| 49 | L | M | 51.6 | 1554.8 | 42 | 1.283 | 1.228 | 0.529 | 59 | 11.416 | 26.353 | 43.702 |
| 50 | L | M | 61.3 | 2091.5 | 42 | 1.592 | 1.460 | 0.496 | 54 | 7.166 | 38.731 | 184.503 |
| 51 | L | M | 16.4 | 421.4 | 14 | 1.258 | 1.173 | 0.452 | 14 | 10.216 | 30.100 | 61.450 |
| 52 | L | F | 0.8 | 441.3 | 1 | 0.767 | 0.767 | 0.000 | 1 | 441.283 | 441.283 | 0.000 |
| 53 | L | M | 5.0 | 1881.6 | 7 | 0.700 | 0.709 | 0.167 | 12 | 37.239 | 156.803 | 282.186 |
| 54 | L | M | 11.2 | 57.2 | 8 | 1.425 | 1.398 | 0.611 | 9 | 6.549 | 6.357 | 3.639 |
| 55 | L | F | 41.0 | 2042.5 | 32 | 1.308 | 1.281 | 0.489 | 37 | 10.449 | 55.202 | 191.986 |
| 56 | L | M | 43.3 | 907.5 | 37 | 1.033 | 1.169 | 0.548 | 47 | 6.166 | 19.308 | 57.095 |
| 57 | L | M | 15.1 | 98.2 | 9 | 1.583 | 1.679 | 0.411 | 11 | 8.773 | 8.927 | 4.511 |
| 58 | L | F | 61.0 | 2125.1 | 49 | 1.200 | 1.245 | 0.540 | 53 | 11.432 | 40.096 | 107.874 |
| 59 | L | M | 17.7 | 713.3 | 17 | 0.917 | 1.043 | 0.378 | 18 | 11.707 | 39.630 | 70.279 |

**Supplementary Table 4.** Duration of locomotion and pause epochs across all the memory and foraging sessions. This table summarizes 59 task-specific epoch sessions (Subjects C, PB, and L) and reports the total time spent (seconds), number of epochs, median, mean, and standard deviation of epoch durations within each session. Tasks abbreviations: F = Foraging, M = Memory alternation, F & M = both tasks within the same session.

| **Session Number** | **Monkey** | **Task** | **Total Time State1 (s)** | **Total Time State2 (s)** | **State1 Epochs** | **State1 Median Epoch Time (s)** | **State1 Mean Epoch Time (s)** | **State1 STD Epoch Time (s)** | **State2 Epochs** | **State2 Median Epoch Time (s)** | **State2 Mean Epoch Time (s)** | **State2 STD Epoch Time (s)** |
| --- | --- | --- | --- | --- | --- | --- | --- | --- | --- | --- | --- | --- |
| 1 | C | F | 1407.7 | 78.7 | 748 | 0.817 | 1.882 | 2.645 | 871 | 0.067 | 0.090 | 0.066 |
| 2 | C | F | 2101.0 | 326.3 | 2684 | 0.458 | 0.783 | 0.906 | 3582 | 0.067 | 0.091 | 0.068 |
| 3 | C | F | 2120.8 | 272.1 | 2281 | 0.533 | 0.930 | 1.185 | 2919 | 0.067 | 0.093 | 0.071 |
| 4 | C | F | 1773.9 | 199.8 | 1708 | 0.550 | 1.039 | 1.699 | 2215 | 0.067 | 0.090 | 0.068 |
| 5 | C | F | 1578.9 | 183.8 | 1644 | 0.533 | 0.960 | 1.187 | 2077 | 0.067 | 0.089 | 0.064 |
| 6 | C | F | 246.0 | 37.0 | 6 | 41.500 | 41.000 | 19.708 | 10 | 3.500 | 3.700 | 1.829 |
| 7 | C | F | 1958.3 | 238.6 | 2174 | 0.500 | 0.901 | 1.230 | 2615 | 0.067 | 0.091 | 0.071 |
| 8 | C | F | 1862.3 | 327.2 | 2429 | 0.483 | 0.767 | 0.879 | 3326 | 0.067 | 0.098 | 0.082 |
| 9 | C | F | 2249.2 | 251.0 | 1884 | 0.583 | 1.194 | 2.182 | 2575 | 0.067 | 0.097 | 0.082 |
| 10 | PB | F | 1847.9 | 94.5 | 876 | 0.725 | 2.109 | 6.480 | 1082 | 0.067 | 0.087 | 0.067 |
| 11 | C | M | 734.3 | 164.6 | 1088 | 0.433 | 0.675 | 0.749 | 1494 | 0.083 | 0.110 | 0.094 |
| 12 | C | M | 1720.1 | 181.1 | 1640 | 0.600 | 1.049 | 1.293 | 2019 | 0.067 | 0.090 | 0.065 |
| 13 | C | M | 2090.5 | 265.2 | 2142 | 0.533 | 0.976 | 1.346 | 2710 | 0.067 | 0.098 | 0.077 |
| 14 | C | M | 1970.2 | 196.3 | 1901 | 0.667 | 1.036 | 1.104 | 2267 | 0.067 | 0.087 | 0.064 |
| 15 | C | M | 2551.0 | 315.1 | 2391 | 0.550 | 1.067 | 1.497 | 3171 | 0.067 | 0.099 | 0.080 |
| 16 | C | M | 3626.2 | 335.5 | 2746 | 0.683 | 1.321 | 1.814 | 3562 | 0.067 | 0.094 | 0.070 |
| 17 | PB | F | 1525.9 | 172.7 | 1911 | 0.483 | 0.798 | 0.916 | 2115 | 0.067 | 0.082 | 0.058 |
| 18 | PB | F | 1320.0 | 98.0 | 1210 | 0.650 | 1.091 | 1.229 | 1240 | 0.067 | 0.079 | 0.055 |
| 19 | PB | F | 842.2 | 61.1 | 731 | 0.617 | 1.152 | 2.015 | 734 | 0.067 | 0.083 | 0.058 |
| 20 | PB | F | 319.9 | 27.7 | 354 | 0.633 | 0.904 | 0.916 | 332 | 0.067 | 0.084 | 0.057 |
| 21 | PB | F | 1164.2 | 88.7 | 1053 | 0.600 | 1.106 | 1.526 | 1062 | 0.067 | 0.084 | 0.060 |
| 22 | PB | F | 1276.2 | 113.7 | 1362 | 0.567 | 0.937 | 1.135 | 1433 | 0.067 | 0.079 | 0.053 |
| 23 | PB | F | 873.0 | 71.5 | 831 | 0.650 | 1.051 | 1.268 | 923 | 0.067 | 0.078 | 0.056 |
| 24 | PB | F | 950.5 | 116.8 | 975 | 0.567 | 0.975 | 1.215 | 1217 | 0.067 | 0.096 | 0.077 |
| 25 | L | F | 57.2 | 7.8 | 52 | 0.467 | 1.099 | 1.449 | 81 | 0.067 | 0.096 | 0.077 |
| 26 | L | M | 289.7 | 25.5 | 219 | 0.717 | 1.323 | 1.578 | 273 | 0.067 | 0.093 | 0.068 |
| 27 | L | F | 60.8 | 4.4 | 38 | 0.758 | 1.601 | 1.755 | 44 | 0.083 | 0.099 | 0.073 |
| 28 | L | M | 1048.7 | 45.8 | 381 | 1.050 | 2.753 | 5.764 | 490 | 0.067 | 0.094 | 0.070 |
| 29 | L | M | 1011.9 | 83.3 | 664 | 0.817 | 1.524 | 2.044 | 852 | 0.067 | 0.098 | 0.074 |
| 30 | L | M | 1449.1 | 119.7 | 883 | 0.783 | 1.641 | 2.696 | 1176 | 0.083 | 0.102 | 0.083 |
| 31 | L | F | 60.3 | 6.4 | 45 | 0.767 | 1.339 | 1.402 | 59 | 0.067 | 0.108 | 0.086 |
| 32 | L | M | 1688.8 | 95.3 | 786 | 0.917 | 2.149 | 4.020 | 991 | 0.083 | 0.096 | 0.071 |
| 33 | L | M | 1679.6 | 118.3 | 919 | 0.917 | 1.828 | 2.756 | 1233 | 0.067 | 0.096 | 0.078 |
| 34 | L | F | 79.0 | 15.6 | 92 | 0.558 | 0.859 | 1.015 | 149 | 0.083 | 0.105 | 0.085 |
| 35 | L | M | 1210.6 | 71.7 | 571 | 0.817 | 2.120 | 4.297 | 794 | 0.067 | 0.090 | 0.070 |
| 36 | L | M | 726.2 | 83.9 | 485 | 0.700 | 1.497 | 2.442 | 751 | 0.083 | 0.112 | 0.099 |
| 37 | L | F | 84.8 | 11.8 | 76 | 0.667 | 1.115 | 1.244 | 109 | 0.083 | 0.108 | 0.092 |
| 38 | L | M | 1881.1 | 147.0 | 1267 | 0.650 | 1.485 | 3.098 | 1558 | 0.067 | 0.094 | 0.071 |
| 39 | L | M | 1749.2 | 65.9 | 561 | 0.833 | 3.118 | 10.396 | 730 | 0.067 | 0.090 | 0.070 |
| 40 | L | F | 276.3 | 33.3 | 6 | 39.500 | 46.047 | 24.882 | 358 | 0.067 | 0.093 | 0.080 |
| 41 | L | M | 417.7 | 25.0 | 9 | 38.000 | 46.413 | 29.475 | 280 | 0.067 | 0.089 | 0.054 |
| 42 | L | M | 396.4 | 24.3 | 215 | 0.750 | 1.844 | 3.450 | 272 | 0.067 | 0.089 | 0.065 |
| 43 | L | F | 217.2 | 40.0 | 256 | 0.567 | 0.849 | 0.765 | 374 | 0.083 | 0.107 | 0.086 |
| 44 | L | M | 733.6 | 67.8 | 468 | 0.842 | 1.567 | 2.705 | 676 | 0.067 | 0.100 | 0.088 |
| 45 | L | M | 1730.9 | 124.8 | 785 | 0.600 | 2.205 | 7.178 | 1183 | 0.083 | 0.105 | 0.085 |
| 46 | L | F | 187.7 | 50.8 | 299 | 0.450 | 0.628 | 0.598 | 482 | 0.067 | 0.105 | 0.092 |
| 47 | L | M | 144.1 | 17.6 | 137 | 0.600 | 1.052 | 1.306 | 182 | 0.067 | 0.097 | 0.075 |
| 48 | L | M | 1368.0 | 141.4 | 911 | 0.583 | 1.502 | 4.585 | 1331 | 0.083 | 0.106 | 0.087 |
| 49 | L | M | 1340.0 | 138.2 | 979 | 0.700 | 1.369 | 2.723 | 1347 | 0.083 | 0.103 | 0.090 |
| 50 | L | M | 1988.8 | 75.7 | 614 | 0.817 | 3.239 | 10.794 | 786 | 0.083 | 0.096 | 0.069 |
| 51 | L | M | 371.8 | 36.4 | 290 | 0.567 | 1.282 | 2.963 | 402 | 0.067 | 0.090 | 0.069 |
| 52 | L | F | 426.7 | 7.0 | 83 | 2.100 | 5.141 | 8.028 | 96 | 0.050 | 0.073 | 0.063 |
| 53 | L | M | 1781.7 | 68.3 | 574 | 0.808 | 3.104 | 7.144 | 770 | 0.067 | 0.089 | 0.081 |
| 54 | L | M | 42.0 | 9.8 | 52 | 0.533 | 0.808 | 0.815 | 98 | 0.083 | 0.100 | 0.066 |
| 55 | L | F | 1905.0 | 94.2 | 803 | 0.800 | 2.372 | 4.724 | 1069 | 0.067 | 0.088 | 0.064 |
| 56 | L | M | 776.9 | 79.5 | 544 | 0.583 | 1.428 | 3.154 | 800 | 0.067 | 0.099 | 0.078 |
| 57 | L | M | 73.0 | 16.8 | 100 | 0.542 | 0.730 | 0.650 | 174 | 0.083 | 0.097 | 0.068 |
| 58 | L | F | 1945.8 | 121.9 | 967 | 0.783 | 2.012 | 3.896 | 1339 | 0.067 | 0.091 | 0.074 |
| 59 | L | M | 629.5 | 55.0 | 418 | 0.767 | 1.506 | 2.283 | 597 | 0.067 | 0.092 | 0.071 |

**Supplementary Table 5.** Duration of state 1 (pause and head stationary) and state 2 (pause and head moving) epochs across all the memory and foraging sessions. This table summarizes 59 task-specific epoch sessions (Subjects C, PB, and L) and reports the total time spent (seconds), number of epochs, median, mean, and standard deviation of epoch durations within each session. Tasks abbreviations: F = Foraging, M = Memory alternation, F & M = both tasks within the same session

| **Session Number** | **Monkey** | **Task** | **SWR Count Locomotion** | **SWR Count Pause** | **SWR Count State 1** | **SWR Count State 2** | **SWR Rate Locomotion** | **SWR Rate Pause** | **SWR Rate State 1** | **SWR Rate State 2** |
| --- | --- | --- | --- | --- | --- | --- | --- | --- | --- | --- |
| 1 | C | F | 0 | 21 | 19 | 2 | 0.000 | 0.014 | 0.013 | 0.025 |
| 2 | C | F | 20 | 139 | 111 | 28 | 0.058 | 0.052 | 0.053 | 0.086 |
| 3 | C | F | 9 | 95 | 86 | 9 | 0.041 | 0.036 | 0.041 | 0.033 |
| 4 | C | F | 1 | 68 | 65 | 1 | 0.008 | 0.033 | 0.037 | 0.005 |
| 5 | C | F | 2 | 22 | 18 | 3 | 0.016 | 0.012 | 0.011 | 0.016 |
| 6 | C | F | 3 | 2 | 2 | 0 | 0.019 | 0.006 | 0.008 | 0.000 |
| 7 | C | F | 10 | 62 | 48 | 5 | 0.028 | 0.026 | 0.025 | 0.021 |
| 8 | C | F | 1 | 9 | 8 | 2 | 0.003 | 0.004 | 0.004 | 0.006 |
| 9 | C | F | 0 | 5 | 5 | 0 | 0.000 | 0.002 | 0.002 | 0.000 |
| 10 | PB | F | 2 | 82 | 82 | 1 | 0.076 | 0.041 | 0.044 | 0.011 |
| 11 | C | M | 36 | 249 | 184 | 51 | 0.235 | 0.227 | 0.251 | 0.310 |
| 12 | C | M | 13 | 318 | 289 | 25 | 0.099 | 0.158 | 0.168 | 0.138 |
| 13 | C | M | 3 | 42 | 30 | 10 | 0.011 | 0.017 | 0.014 | 0.038 |
| 14 | C | M | 2 | 14 | 13 | 1 | 0.021 | 0.006 | 0.007 | 0.005 |
| 15 | C | M | 60 | 394 | 357 | 44 | 0.179 | 0.132 | 0.140 | 0.140 |
| 16 | C | M | 7 | 118 | 110 | 10 | 0.023 | 0.029 | 0.030 | 0.030 |
| 17 | PB | F | 29 | 166 | 158 | 10 | 0.136 | 0.087 | 0.104 | 0.058 |
| 18 | PB | F | 0 | 380 | 329 | 15 | 0.000 | 0.240 | 0.249 | 0.153 |
| 19 | PB | F | 26 | 162 | 147 | 17 | 0.809 | 0.158 | 0.175 | 0.278 |
| 20 | PB | F | 0 | 92 | 85 | 3 | 0.000 | 0.214 | 0.266 | 0.108 |
| 21 | PB | F | 1 | 6 | 6 | 0 | 0.026 | 0.004 | 0.005 | 0.000 |
| 22 | PB | F | 17 | 150 | 134 | 6 | 0.265 | 0.096 | 0.105 | 0.053 |
| 23 | PB | F | 1 | 99 | 85 | 14 | 0.111 | 0.096 | 0.097 | 0.196 |
| 24 | PB | F | 0 | 6 | 6 | 0 | 0.000 | 0.005 | 0.006 | 0.000 |
| 25 | L | F | 0 | 0 | 0 | 0 | 0.000 | 0.000 | 0.000 | 0.000 |
| 26 | L | M | 0 | 86 | 80 | 6 | 0.000 | 0.267 | 0.276 | 0.235 |
| 27 | L | F | 1 | 2 | 3 | 0 | 0.219 | 0.030 | 0.049 | 0.000 |
| 28 | L | M | 0 | 58 | 56 | 2 | 0.000 | 0.053 | 0.053 | 0.044 |
| 29 | L | M | 29 | 461 | 445 | 21 | 0.671 | 0.412 | 0.440 | 0.252 |
| 30 | L | M | 8 | 199 | 186 | 11 | 0.121 | 0.124 | 0.128 | 0.092 |
| 31 | L | F | 1 | 96 | 91 | 5 | 0.172 | 1.430 | 1.510 | 0.785 |
| 32 | L | M | 17 | 773 | 735 | 36 | 0.843 | 0.429 | 0.435 | 0.378 |
| 33 | L | M | 0 | 749 | 687 | 55 | 0.000 | 0.412 | 0.409 | 0.465 |
| 34 | L | F | 0 | 7 | 7 | 0 | 0.000 | 0.073 | 0.089 | 0.000 |
| 35 | L | M | 0 | 139 | 130 | 8 | 0.000 | 0.107 | 0.107 | 0.112 |
| 36 | L | M | 0 | 17 | 14 | 3 | 0.000 | 0.020 | 0.019 | 0.036 |
| 37 | L | F | 0 | 0 | 0 | 0 | 0.000 | 0.000 | 0.000 | 0.000 |
| 38 | L | M | 3 | 107 | 96 | 11 | 0.026 | 0.051 | 0.051 | 0.075 |
| 39 | L | M | 7 | 876 | 847 | 28 | 0.443 | 0.476 | 0.484 | 0.425 |
| 40 | L | F | 3 | 743 | 657 | 43 | 0.592 | 2.387 | 2.378 | 1.292 |
| 41 | L | M | 0 | 575 | 558 | 4 | 0.000 | 1.252 | 1.336 | 0.160 |
| 42 | L | M | 1 | 213 | 197 | 14 | 0.408 | 0.494 | 0.497 | 0.577 |
| 43 | L | F | 0 | 7 | 6 | 0 | 0.000 | 0.025 | 0.028 | 0.000 |
| 44 | L | M | 0 | 80 | 74 | 6 | 0.000 | 0.097 | 0.101 | 0.089 |
| 45 | L | M | 13 | 464 | 419 | 33 | 0.185 | 0.242 | 0.242 | 0.264 |
| 46 | L | F | 2 | 22 | 20 | 3 | 0.067 | 0.084 | 0.107 | 0.059 |
| 47 | L | M | 1 | 25 | 20 | 3 | 0.054 | 0.144 | 0.139 | 0.171 |
| 48 | L | M | 33 | 349 | 312 | 34 | 0.412 | 0.221 | 0.228 | 0.240 |
| 49 | L | M | 1 | 69 | 65 | 4 | 0.019 | 0.044 | 0.049 | 0.029 |
| 50 | L | M | 3 | 45 | 42 | 3 | 0.049 | 0.022 | 0.021 | 0.040 |
| 51 | L | M | 4 | 94 | 85 | 11 | 0.244 | 0.223 | 0.229 | 0.302 |
| 52 | L | F | 0 | 37 | 37 | 0 | 0.000 | 0.084 | 0.087 | 0.000 |
| 53 | L | M | 1 | 826 | 785 | 35 | 0.201 | 0.439 | 0.441 | 0.512 |
| 54 | L | M | 2 | 5 | 4 | 1 | 0.179 | 0.087 | 0.095 | 0.102 |
| 55 | L | F | 9 | 60 | 55 | 5 | 0.220 | 0.029 | 0.029 | 0.053 |
| 56 | L | M | 6 | 27 | 30 | 0 | 0.139 | 0.030 | 0.039 | 0.000 |
| 57 | L | M | 3 | 22 | 21 | 2 | 0.198 | 0.224 | 0.287 | 0.119 |
| 58 | L | F | 17 | 619 | 596 | 27 | 0.279 | 0.291 | 0.306 | 0.222 |
| 59 | L | M | 14 | 369 | 339 | 27 | 0.790 | 0.517 | 0.539 | 0.491 |

**Supplementary Table 6.** SWR rates across states. This table reports the SWR counts and SWR rates (events/s) computed separately for the movement states. Columns include session number, subject, task (F = Foraging, M = Memory alternation), number of SWRs in locomotion, pauses, state 1 and state 2, and the corresponding SWR rate (events/s), computed as number of SWRs divided by total time spent in that state for that session
